# Enhancer buffering protects dosage-sensitive housekeeping genes during vulnerable developmental transitions

**DOI:** 10.64898/2026.07.29.741571

**Authors:** Chunyang Ni, Lucia Ichino, Tomek Swigut, Joanna Wysocka

## Abstract

Housekeeping genes maintain robust expression across cell types despite dynamic transcription factor fluctuations, yet their haploinsufficiency is associated with many tissue-specific developmental disorders. To understand this paradox, we focus on TCOF1, a broadly expressed regulator of rRNA synthesis, whose haploinsufficiency causes Treacher Collins syndrome (TCS). We show that transitional cranial neural crest cells (tCNCC) undergoing mesenchymal specification exhibit extreme dosage-sensitivity to TCOF1, but not to RNA polymerase I, explaining both cellular origins of TCS and prevalence of TCOF1 mutations in the disease. To maintain robust expression, TCOF1 deploys CNCC-specific enhancers that buffer against fluctuations in promoter-regulating factors, converting sensitive expression responses into threshold-protected outputs. Systematic promoter-enhancer coupling experiments demonstrate this principle generalizes across many dosage-sensitive housekeeping genes, whose promoters operate near saturation and only reveal their enhancer-dependency under suboptimal conditions. Thus, enhancers play a unique role at housekeeping promoters: rather than amplifying expression, they ensure its robustness during vulnerable developmental transitions.

## Main Text

Housekeeping genes are broadly expressed across cell types and associated with maintenance of basic cellular functions, such as cell growth, ribosome biogenesis, proteostasis and metabolism. In contrast to developmental genes, housekeeping gene expression is generally less dependent on cell type-specific enhancers (1–6). Instead, housekeeping gene transcription is driven by extended promoters and promoter-proximal sequences, which are regulated by ubiquitously expressed transcription factors and can sustain robust expression levels across many cell types (1–5, 7, 8).

Despite their broad and robust expression, housekeeping genes often exhibit dosage sensitivity and their haploinsufficiency is associated with a wide range of human congenital diseases characterized by tissue-specific manifestations. Indeed, recent analyses of copy number variation patterns in nearly one million human genomes predict that nearly 3000 human genes are haploinsufficient (*9*); many of these dosage-sensitive genes show broad expression patterns and essentiality across many cell types. For example, *TCOF1* is a housekeeping gene encoding a protein involved in rRNA synthesis by RNA polymerase I and organization of the fibrillar center in the nucleolus, and thus critical for ribosome biogenesis (*10–12*). *TCOF1* haploinsufficiency causes Treacher Collins Syndrome (TCS), a congenital craniofacial disorder characterized by mandibulofacial dysostosis (*13–16*). Despite a broad role of TCOF1 (also referred to as Treacle) in ribosome biogenesis, craniofacial manifestations of TCS occur in the absence of defects in other organs, and have been linked to the cranial neural crest, an embryonic cell population that gives rise to the majority of the facial structures (*17–21*). Consistently with rRNA synthesis defects being the root cause of the disease, TCS or TCS-like mandibulofacial dysostosis can also result from heterozygous mutations in genes encoding RNA polymerase I subunits (*POLR1A*, *POLR1B*, *POLR1C*, *POLR1D*) (*22–26*). However, *TCOF1* dominates the mutagenic spectrum of TCS, with 85-90% of TCS cases being attributable to the *TCOF1* loss-of-function mutations (*15, 16*). Notably, this haploinsufficiency pattern cannot be explained by differences in gene or protein size, as *POLR1A* and *POLR1B* have comparable genomic spans or encode proteins of similar molecular weight to TCOF1, yet show fundamentally different mutational tolerances in human disease. These observations raise several questions: What underlies cell type-selective vulnerability to changes in TCOF1 dosage? What is the molecular basis for *TCOF1*’s genetic prevalence over *POLR1* genes, despite their functional convergence on the same biochemical pathway? How is TCOF1 dosage stabilized during vulnerable cell fate transitions to facilitate normal development?

To address these questions, we combine human cranial neural crest cell (CNCC) differentiation models with tunable protein dosage control. We identify transitional CNCC states undergoing mesenchymal fate commitment as the precise developmental window of vulnerability to TCOF1 dosage reduction. We further explain genetic prevalence of *TCOF1* mutations in TCS by demonstrating that TCOF1 dosage is highly limiting for rRNA synthesis and differentiation of CNCC, whereas POLR1A/B exhibit robust buffering capacity beyond a two-fold reduction in levels. In addition, CNCC show intrinsic hypersensitivity to rRNA synthesis perturbation and this compounded vulnerability—TCOF1’s dosage hypersensitivity coupled with CNCC-specific stress sensitivity—renders TCOF1 dosage critical for CNCC development.

The extreme dosage sensitivity, coupled with successful CNCC development in normal individuals, points to the existence of regulatory mechanisms that maintain robust control of TCOF1 expression during sensitive cell fate transitions. We identify a set of CNCC developmental enhancers that support TCOF1 expression, safeguarding mesenchymal fate commitment. However, rather than amplifying TCOF1 levels, these enhancers function as dosage buffers against transient fluctuations in promoter regulator MYC. Further systematic promoter-enhancer coupling experiments demonstrate that this buffering mechanism generalizes across dosage-sensitive housekeeping genes with a broad range of expression. We propose that the role of cell type-specific enhancers in regulation of developmental and housekeeping genes is fundamentally distinct: while enhancers often amplify expression at developmental promoters, at dosage-sensitive housekeeping genes they convert sensitive responses to transcription factor fluctuations into buffered, threshold-protected outputs, ensuring expression robustness during vulnerable developmental transitions.

### In vitro differentiation model captures transitional states during human CNCC specification

TCS has been widely recognized as a neurocristopathy, but the precise developmental transition where the disease manifests has remained elusive due to a lack of relevant human models, although previous work in mice suggests that it originates in the neuroepithelium (*14*). We have previously established an in vitro differentiation model for derivation of cranial neural crest cells (CNCC) from human embryonic stem cells (hESC) (*27, 28*). In this model, ESC are first induced to form neuroepithelial spheres that subsequently attach and give rise to delaminating CNCC that undergo epithelial-to-mesenchymal (EMT) transition and migrate away. These migratory cells are then isolated after 11 days of differentiation for the long term maintenance of homogenous CNCC population with mesenchymal characteristics (*28*). We hypothesized that cell population at an earlier differentiation time point (day 8), which contains both attached neuroepithelial spheres and delaminating CNCC, could capture transient cellular states relevant for the TCS etiology (Fig. 1A). To synchronize differentiation and enable accurate capture of transient developmental trajectories, we optimized the protocol by using Aggrewell 800 plates to seed uniform ESC numbers for initial neuroepithelial sphere formation (see Methods). This ensured highly reproducible neural induction, sphere attachment and delamination across replicates and enabled robust attribution of observed cellular states to developmental progression (Fig. S1G). To investigate cellular heterogeneity of the day 8 cell population, we profiled these cells using single cell RNA-seq (scRNA-seq) and performed unsupervised clustering of the resulting data. This analysis revealed multiple cell clusters expressing markers consistent with cell fate transitions occurring during CNCC formation, as previously described in model organisms (*17, 18, 29*), including: dorsal neural tube (NT), pre-EMT neural crest (pre-EMT NC), and three intermediate populations positioned between pre-EMT neural crest and fully committed mesenchymal cells, which we termed transitional CNCC (tCNCC) (Fig. 1B,C; Fig. S1A,B) (*29*). Additional clusters representing pluripotent/early neuroectoderm and neuronal cells were also detected as expected byproducts of differentiation (Fig. S1A,B). This cellular heterogeneity mirrors cellular diversity previously described in the scRNA-seq analysis of the mouse embryonic neural crest development, suggesting that our human *in vitro* system recapitulates many aspects of the embryonic neural crest specification (*29*). The molecular signatures of the three tCNCC sub-clusters are consistent with the stepwise fate specification and acquisition of the mesenchymal potential, with the tCNCC1 characterized by high expression of SOX10 and FOXD3, marking early neural crest identity, and tCNCC2 and tCNCC3 progressively upregulating TWIST1, a major transcription factor in the regulation of neural crest mesenchyme (*18, 30*) (Fig. 1C). In addition, tCNCC1 and tCNCC2 share expression of the surface marker ITGA4 (CD49D), linking early specification and transitional states, and tCNCC3 expresses surface marker MME (CD10), consistent with a more mature mesenchymal state; we will later rely on these surface markers to enrich the relevant cell populations (Fig. S1C,D) (*28*). Pseudotime analysis confirmed an ordered developmental trajectory from pre-EMT cells through tCNCC1>2>3 (Fig. 1D), indicating that CNCC specification in vitro proceeds through discrete transitional states resembling those previously described in mouse and avian embryos (*17, 18, 29, 31*). In humans, CNCC specification and delamination occurs within 2-4 weeks of gestation – the least accessible window in human embryology which extends beyond the 14 day timepoint when the human embryos can be cultured in vitro but occurs before pregnancies are recognized. Thus, the stem cell-based model described here provides a valuable tool for studies of early CNCC cell fate decisions and modeling neurocristopathies.

**Figure 1.**
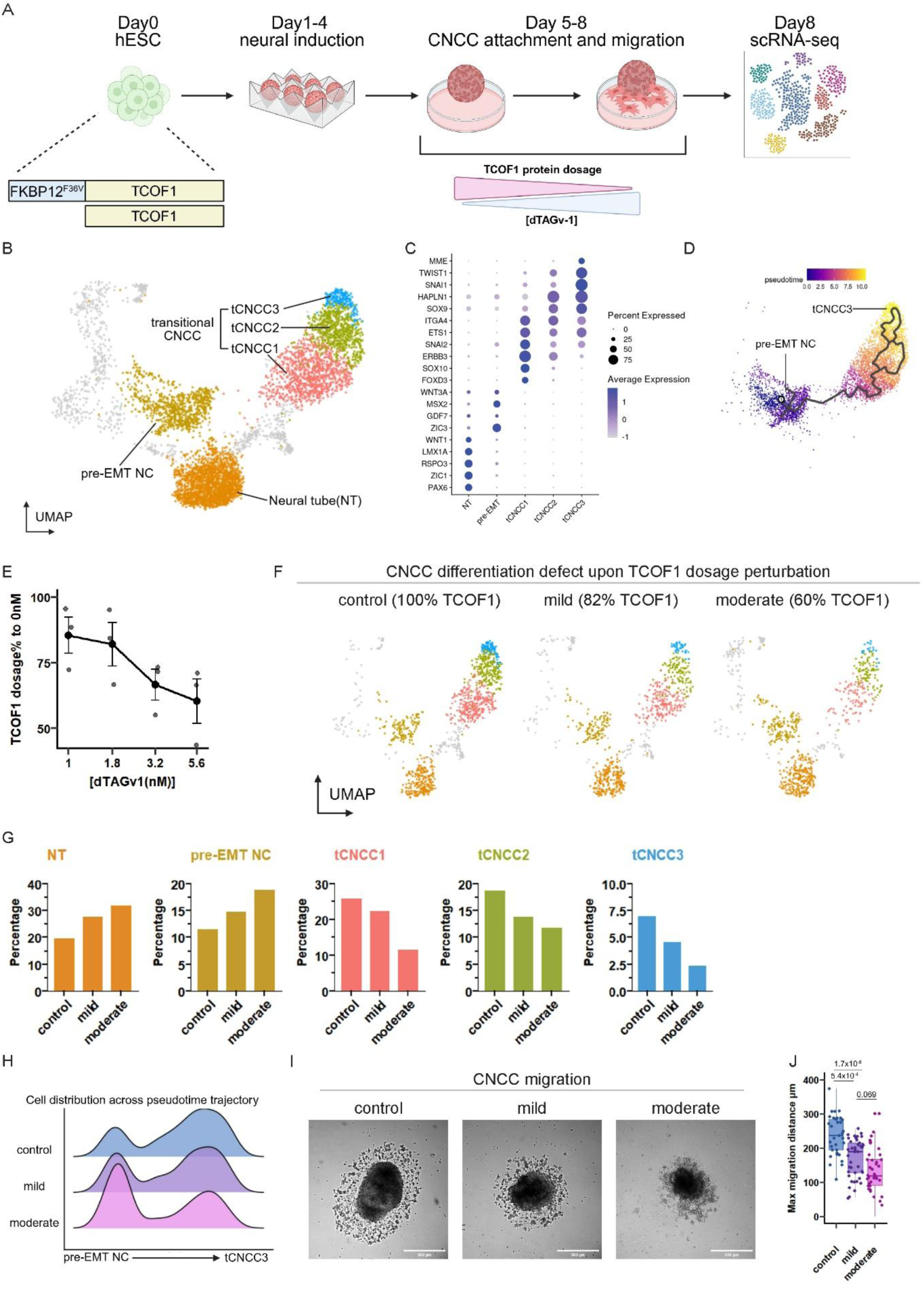
Transitional cranial neural crest cell states are hypersensitive to TCOF1 dosage. **(A)** Schematic overview of cranial neural crest cell (CNCC) differentiation model with TCOF1 degron-mediated dosage control and single-cell RNA-seq profiling at day 8. **(B)** UMAP embedding of day 8 cells reveals discrete populations spanning neural tube (NT), pre-EMT neural crest to three sequential transitional CNCC states (tCNCC1-3) as highlighted. **(C)** Expression of key developmental markers across the NT to CNCC trajectory. **(D)** Pseudotime trajectory shows developmental progression from pre-EMT neural crest through tCNCC1→tCNCC2→tCNCC3. **(E)** Flow cytometry quantification of TCOF1 protein levels in combined tCNCC populations (tCNCC1-3) following dTAGv-1 treatment at different concentrations, shown as % relative to same cells treated with vehicle without dTAGv-1. Mild depletion (1.8 nM): 82% of control; moderate depletion (5.6 nM): 60% of control. Error bars represent SEM, from n=3 independent differentiations. **(F)** UMAP embedding split by TCOF1 dosage showing progressive loss of tCNCC populations. **(G)** Quantification of cell type proportions across TCOF1 dosage. TCOF1 reduction causes dose-dependent depletion of tCNCC populations, while NT and pre-EMT NC increase their relative proportion in the population. Data from one representative replicate (n=2 independent differentiations). **(H)** Distribution of cell populations across the pseudotime trajectory shows TCOF1-depleted cells are defective in progression from pre-EMT NC to later stages. **(I)** Representative images of day 7 cultures showing impaired cell migration from neuroepithelial spheres with TCOF1 depletion. Scale bars, 500 μm. **(J)** Quantification of migration distance from neuroepithelial spheres. Boxplot from n=33, 41, 38 spheres across 3 independent differentiations. Error bars represent SEM. Kruskal-Wallis test followed by Dunn’s post-hoc test with Bonferroni adjustment.

### High vulnerability of transitional CNCC states to changes in TCOF1 dosage

Having established a model system capable of capturing transient CNCC developmental states, we next sought to identify which specific cellular transition(s) are most vulnerable to *TCOF1* haploinsufficiency. To model TCS in our human differentiation system, we engineered a heterozygous FKBP12^F36V^ degron tag at the endogenous *TCOF1* locus in ESC. In this setting, degron activation by addition of the dTAGv-1 molecule can decrease TCOF1 protein levels by up to two-fold, mimicking haploinsufficiency in TCS patients (Fig. 1A) (*32, 33*). By applying distinct dTAGv-1 concentrations we can achieve distinct degree of TCOF1 dose perturbation, as measured by the intracellular staining with a TCOF1 antibody followed by flow cytometry (Fig. 1E). Subsequently, we focused on the 1.8 nM or 5.6 nM, concentrations, which resulted in, respectively, mild (82% of the endogenous TCOF1 dosage) or moderate (60% of the endogenous TCOF1 dosage) reduction in protein levels. Analysis of the resulting cell populations at day 8 of differentiation by scRNA-seq revealed TCOF1 dose-dependent, progressive decrease of the tCNCC populations, whereas the NT or pre-EMT NC populations were unaffected or even increased in their relative proportion (Fig. 1F,G, Fig. S1E,F). The reduction in tCNCC yield was evident even upon mild depletion, indicating that downregulation of TCOF1 levels by 20% is sufficient to alter CNCC differentiation trajectory. Pseudotime analysis is consistent with TCOF1 reduction leading to a developmental failure, with cells struggling to progress from pre-EMT through later tCNCC stages (Fig. 1H). In agreement with the strong effect on the mesenchymal tCNCC2/3 population, TCOF1-reduced cultures also showed dose-dependent impaired migration capacity, with death of the delaminating cells evident at the moderate depletion levels (Fig. 1I,J). Together, these results identify high vulnerability of the transitory CNCC states to perturbations in TCOF1 dosage.

### TCOF1 but not POLR1 shows high dosage sensitivity in transitional CNCC

Heterozygous *TCOF1* mutations account for a vast majority (approximately 85-90%) of known TCS-spectrum cases (*15, 16, 34, 35*), distribute across the entire gene length without apparent hotspots and include cases with complete single-copy loss, suggesting haploinsufficiency as the primary disease mechanism (Fig. 2A) (*36*). In contrast, reported mutations in *POLR1A/B/C/D* are predominantly missense variants (*22–25*), with complete haploinsufficiency rarely documented, raising the possibility of differential dosage tolerance among TCS genes.

**Figure 2.**
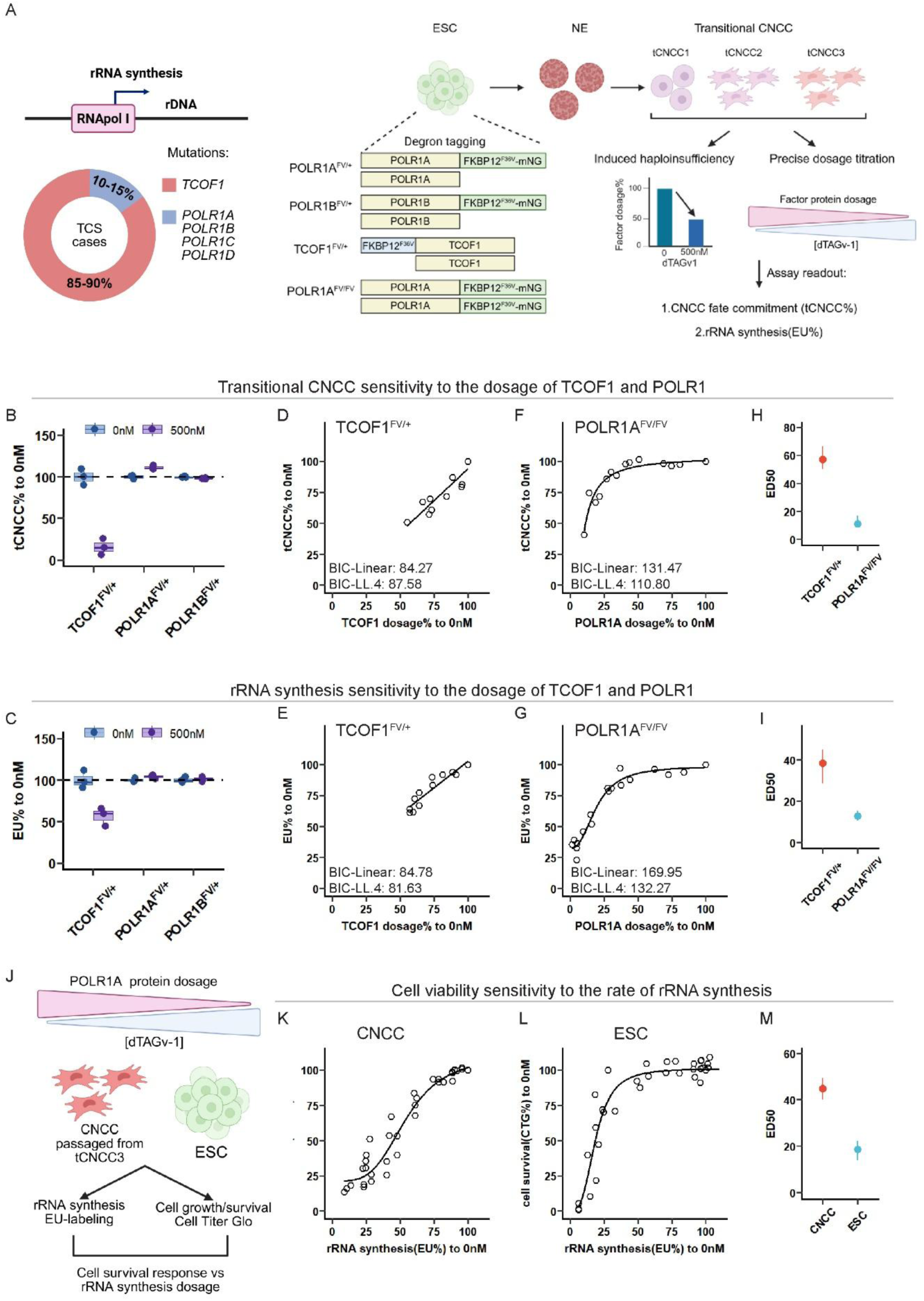
TCOF1 exhibits unique dosage sensitivity compared to POLR1 components. **(A)** Left, schematic showing that *TCOF1* mutations dominate Treacher Collins syndrome genetics (85-90% of cases). Right, degron tagging strategy for induced haploinsufficiency and quantitative protein titration. **(B)** tCNCC yield (measured by surface marker staining and flow cytometry) following induced haploinsufficiency (96 hours dTAGv-1 treatment) of TCOF1, POLR1A, or POLR1B, quantified as % relative to vehicle treated control cells. n=3 independent differentiations. Statistical comparisons control(0nM) vs dTAGv-1(500nM): TCOF1 (p = 1.5×10-3, mean difference= -84.3%), POLR1A (p = 0.015, mean difference= 11.2%, higher in treated condition), POLR1B (p = 0.059, mean difference= -1.97%), two sided t-test with bonferroni correction. **(C)** Ribosomal RNA synthesis (5-EU incorporation) following induced haploinsufficiency (96 hours dTAGv-1 treatment) of TCOF1, POLR1A, or POLR1B in tCNCC3, quantified as % relative to vehicle treated control cells. n=3 independent differentiations. Statistical comparisons control(0nM) vs dTAGv-1(500nM): TCOF1 (p = 0.023, mean difference= -43.4%), POLR1A (p = 0.26), POLR1B (p = 1.0), two sided t-test with bonferroni correction. **(D)** tCNCC yield across TCOF1 dosage titration shows linear dose-response, as indicated by lower Bayesian information criterion (BIC) for the linear model compared to alternative four-parameter log-logistic model (LL4). TCOF1 protein levels were measured at the 96h endpoint following dTAGv-1 treatment as percentage relative to vehicle (0nM) control. n=3 independent differentiations. **(E)** Ribosomal RNA synthesis (5-EU) in tCNCC3 across TCOF1 dosage titration shows linear dose-response. Linear model is preferred over LL4 based on comparable BIC values and model parsimony. TCOF1 protein levels were measured at the 24h endpoint following dTAGv-1 treatment as percentage relative to vehicle (0nM) control. n=2 independent differentiations. **(F)** tCNCC yield across POLR1A dosage shows buffered response, with better fit (lower BIC) with the non-linear LL4 model. POLR1A protein levels were measured at the 96h endpoint following dTAGv-1 treatment as percentage relative to vehicle (0nM) control. n=3 independent differentiations. **(G)** Ribosomal RNA synthesis (5-EU) in tCNCC3 across POLR1A dosage shows buffered response, with better fit (lower BIC) with the non-linear LL4 model. POLR1A protein levels were measured at the 24h endpoint following dTAGv-1 treatment as percentage relative to vehicle (0nM) control. n=3 independent differentiations. **(H)** ED50 value comparison for tCNCC yield (predicted protein dosage corresponding to 50% tCNCC yield) between TCOF1 and POLR1A. 95% confidence intervals were obtained by bootstrap resampling of individual data points (2,000 iterations), with model refitting and ED estimation repeated per iteration. **(I)** ED50 value comparison for rRNA synthesis (predicted protein dosage corresponding to 50% EU signal relative to control) between TCOF1 and POLR1A. 95% confidence intervals were obtained by bootstrap resampling of individual data points (2,000 iterations), with model refitting and ED estimation repeated per iteration. **(J)** Schematic of experimental design for testing cell type-specific sensitivity to rRNA synthesis: matched rRNA synthesis perturbations in CNCC and ESC were examined along with measurements of cell viability. **(K)** CNCC viability (measured by CellTiter-Glo luminescence relative to no treatment control) shows steep decline with rRNA synthesis reduction, fit with the non-linear LL4 model. rRNA synthesis (EU) and cell survival (CTG) levels were measured at the 24h endpoint following dTAGv-1 treatment as percentage relative to vehicle (0nM) control, n=5 independent differentiations. **(L)** ESC viability (measured by CellTiter-Glo luminescence relative to no treatment control) shows buffered response to comparable rRNA synthesis reduction, fit with the non-linear LL4 model. rRNA synthesis (EU) and cell survival (CTG) levels were measured at the 24h endpoint following dTAGv-1 treatment as percentage relative to vehicle (0nM) control, n=4 independent differentiations. **(M)** ED50 value comparison for cell viability (predicted cell viability corresponding to 50% reduction in rRNA synthesis relative to control) for CNCC and ESC. 95% confidence intervals were obtained by bootstrap resampling of individual data points (2,000 iterations), with model refitting and ED estimation repeated per iteration.

To determine whether *TCOF1*’s genetic prevalence reflects unique dosage sensitivity, we generated heterozygous FKBP12^F36V^-mNG tagged ESC lines for POLR1A and POLR1B, enabling matched quantitative depletion within 100% to 50% of physiological dosage range, along with monitoring POLR1A/B levels by mNG fluorescence (Fig. 2A). In addition, we also generated homozygously FKBP12^F36V^-mNG degron-tagged ESC lines for POLR1A, allowing for acute and precise perturbation of POLR1A dosage across the entire physiological range (Fig. 2A). We first used a set of heterozygous TCOF1, POLR1A or POLR1B ESC to selectively deplete the tagged protein product corresponding to one of the two alleles (which we termed ‘induced haploinsufficiency’); see Fig. S2A for validation of nearly complete loss of the tagged proteins upon addition of the degron-inducing dTAGv-1 molecule. We then compared CNCC differentiation efficiency in the induced haploinsufficiency conditions to the same cells without addition of the dTAGv-1 by quantifying tCNCC population (tCNCC1/2/3) at day 8 of differentiation by flow cytometry (Fig. 2B; Fig. S2B). In parallel, we measured rRNA synthesis rates in day 8 tCNCC3 population by pulse-labeling with 5-ethynyl uridine (5-EU), followed by click chemistry conjugation to Azide-568 and flow cytometry quantification of EU incorporation. We observed that only TCOF1 induced haploinsufficiency impaired tCNCC differentiation and rRNA synthesis, whereas reducing the dose of POLR1A and POLR1B by ∼50% does not impair either tCNCC yield or rRNA synthesis (Fig. 2B,C, Fig. S2C).

To quantitatively characterize dosage responses, we performed systematic dTAGv-1 titrations spanning 100% to 50% dosage for TCOF1 and 100% to near-complete depletion for POLR1A. TCOF1 exhibited a steep, linear dose-response curve: almost every incremental loss of TCOF1 reduced both tCNCC yield and rRNA synthesis (Fig. 2D, E). POLR1A, by contrast, demonstrated robust buffering capacity—both tCNCC differentiation and rRNA synthesis remained nearly unperturbed until the POLR1A dose reached below 40% of the endogenous level, beyond which both collapsed sharply (Fig. 2F,G). We next calculated the median effective dose (ED50) of a fitted linear model for TCOF1 and log-logistic model for POLR1A dose response curves, corresponding to the protein dose at which tCNCC yield (Fig. 2H) or rRNA synthesis (Fig. 2I) is reduced by half. Consistent with pronounced dosage sensitivity, these quantifications revealed much higher ED50 for TCOF1 in both responses, as compared to POLR1A (Fig. 2H,I). For example, the tCNCC yield is reduced by half when TCOF1 dosage reaches 57% of the endogenous level; in contrast the same yield reduction only occurs when POLR1A levels are diminished to 11% of the endogenous level. Thus, while both TCOF1 and POLR1A are required for rRNA synthesis and CNCC differentiation, TCOF1 dosage is much more limiting in these processes, whereas POLR1A exhibits strong buffering until severe disruption beyond the 50% reduction. This differential sensitivity resolves the longstanding question of why heterozygous *TCOF1* mutations dominate TCS genetics: subtle perturbations to TCOF1 protein levels trigger developmental defects while similar perturbations to POLR1 are well-tolerated by cells.

### CNCC are highly sensitive to perturbations in rRNA synthesis

TCOF1’s dosage sensitivity explains its genetic prevalence in TCS but it leaves unresolved the question of why CNCC are uniquely sensitive to perturbations in rRNA synthesis, in turn leading to highly tissue-specific manifestations of the disease. Previous work provided one potential explanation by suggesting that CNCC show heightened sensitivity to p53-mediated nucleolar stress response and apoptosis, which is also consistent with observations that inactivation of p53 partially rescues TCS craniofacial manifestations in mice (*20, 21*). Our POLR1A titration system offers a model to quantitatively compare effects of rRNA synthesis on cell viability between CNCC and other cell types. To this end, we titrated POLR1A levels in parallel CNCC and ESC cultures to achieve matched reductions in rRNA synthesis, as measured by EU incorporation (Fig. 2J). After 24h of dTAGv-1 treatment, we quantified effects of these reductions on cell viability, assayed by luminescence (CellTiter-Glo). Across matched levels of rRNA synthesis reduction, CNCC showed steep viability decline even at modest rRNA perturbation, whereas ESC tolerated substantial rRNA reduction with minimal impact (Fig. 2K,L). This differential sensitivity was reflected in differences in ED50 of the viability responses to changes in rRNA synthesis between CNCC and ESC (Fig. 2M). We then performed RNA-seq on CNCC and ESC under matched titration conditions, achieving comparable rRNA synthesis reduction kinetics (6 h: ∼70%, 12 h: ∼50%, 24 h: ∼30% 5-EU signal compared to unperturbed control) (Fig. S3A-B). The results confirmed the underlying molecular basis: CNCC rapidly upregulated genes associated with p53-associated stress responses. This upregulation was already evident at 6 h (when rRNA synthesis was only modestly reduced) and intensified to a strong response by 24 h. In contrast, ESC showed delayed and weaker transcriptional responses, with no response at 6 h and a relatively weak response at 24 h (Fig. S3C,D). Together, our results suggest that this cellular hypersensitivity to rRNA synthesis levels, when combined with TCOF1’s dosage sensitivity, may create a compounded vulnerability, providing an explanation for both the prevalence of the *TCOF1* heterozygosity and tissue selectivity of TCS.

### Developmental enhancers support stage-specific *TCOF1* expression

*TCOF1* is broadly expressed across tissues and regulated by a GC-rich promoter (*36*). Typically, such housekeeping gene promoters can sustain high expression of their target genes without relying on distal enhancer activity (*1*). However, given the extreme TCOF1 dosage-sensitivity in CNCC, we hypothesized that additional cis-regulatory elements beyond the promoter may support its expression during CNCC formation to facilitate robustness of the developmental process. To identify such candidate elements, we used FACS with panel of surface markers described in Figure S1C to isolate neuroepithelial (NE) and three transitional CNCC populations (tCNCC1-3) at day 8 of CNCC differentiation and performed bulk ATAC-seq analysis for each of these populations. We further included the ATAC-seq data and epigenomic profiles from ESC and mesenchymal CNCC of the matched genetic background from our previously published studies (*28, 37*). Inspection of the chromatin profiles at the *TCOF1* locus revealed several promoter-distal open chromatin peaks (Fig. 3A). One of the peaks was constitutively open across cell types and overlapped with a CTCF and cohesin (RAD21) binding site in CNCC, suggesting a potential structural role. However, three other distal open chromatin regions were dynamically regulated and enriched for H3K27ac in CNCC, consistent with a potential role as enhancers: E1 (+20 kb), E2 (−37 kb), and E3 (−50 kb) from the *TCOF1* promoter (Fig. 3A). These candidate enhancers were inaccessible in ESC, gained accessibility during differentiation, especially at the tCNCC2 and tCNCC3 stages, and remained open in CNCC (Fig. 3A). To confirm their regulatory potential, we cloned the E1-E3 regions, separately or together, as well as the region harboring the CTCF peak, and tested their activity in the luciferase reporter assay in CNCC, along with the strong SV40 viral enhancer as a positive control. All three candidate enhancers, but not the CTCF-bound region, exhibited autonomous regulatory activity, with the E1 and E2 exceeding the activity of the SV40 viral enhancer, and the E123 compound enhancer showing strong, synergistic activity (Fig. 3B). The opening of the enhancers at the *TCOF1* locus in the tCNCC2 and tCNCC3 populations coincides with the acquisition of the mesenchymal states including expression of the key migratory/mesenchymal fate regulator TWIST1 (*17, 18, 30, 31*). Notably, our previously published work reveals TWIST1 is a central transcriptional regulator in mesenchymal CNCC, required for chromatin opening at thousands of enhancer elements (*38*). Examination of our previously reported ChIP-seq data revealed that TWIST1 indeed binds at E1-E3 enhancers in CNCC (Fig. 3D) (*38*). Moreover, acute degradation of TWIST1 in CNCC results in decreased chromatin accessibility and H3K27ac at these regions, confirming functional regulation of E1-E3 enhancers by TWIST1 (Fig. 3C,D) (*38*).

**Figure 3.**
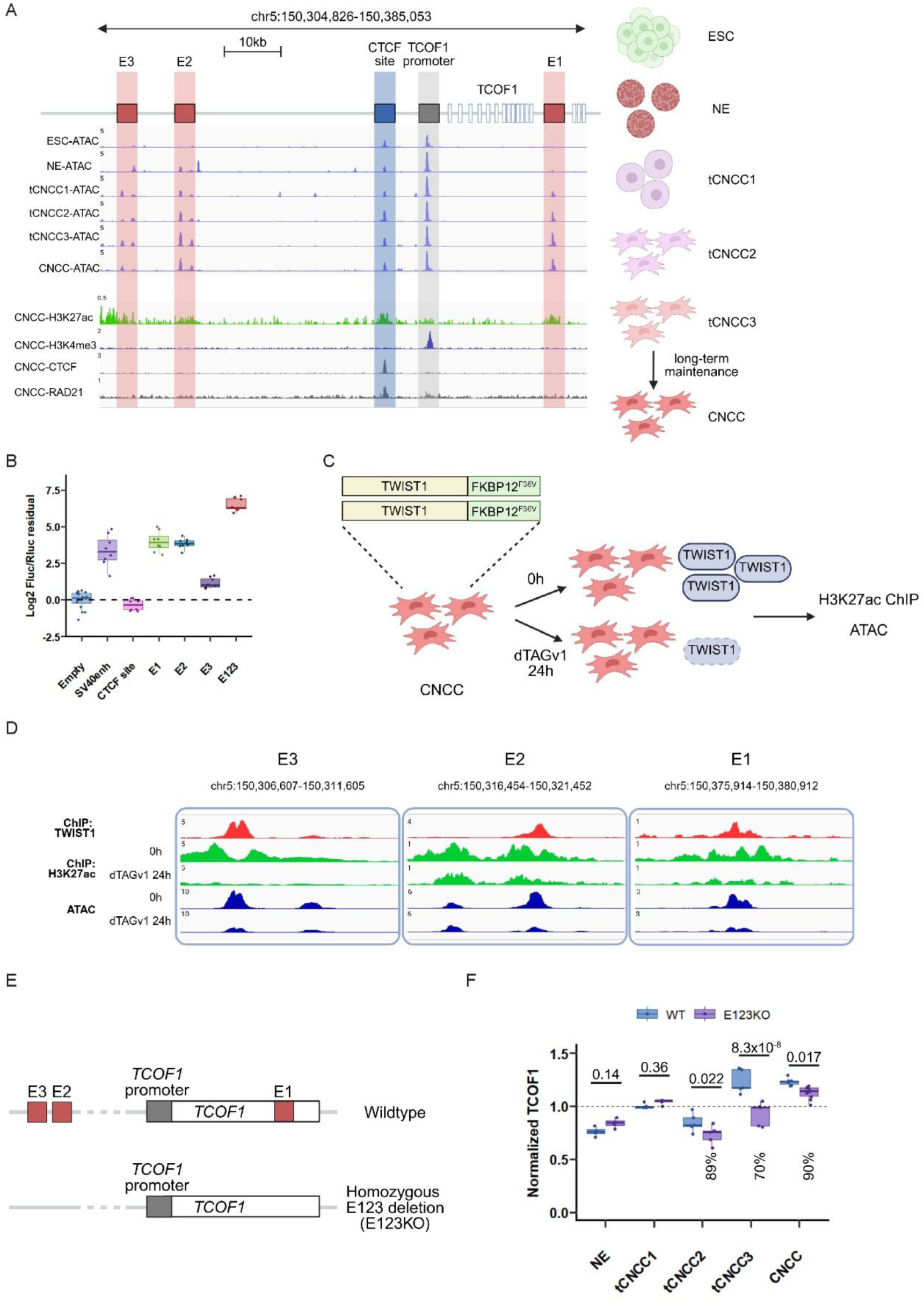
Developmental enhancers sustain *TCOF1* expression during CNCC mesenchymal transition. **(A)** Chromatin accessibility (ATAC-seq) tracks at the *TCOF1* locus are shown across differentiation stages. Data from one representative replicate (n=2 independent differentiations; see deposit data). In addition, ChIP-seq tracks for histone marks (H3K27ac, H3K4me3), CTCF and RAD21 occupancy from genetically matched long-term maintained CNCC are shown at the bottom. Three candidate enhancers (E1, E2, E3), CTCF/RAD21 site and promoter are schematically depicted on top. **(B)** Luciferase reporter assay of individual enhancers (E1, E2, E3), CTCF site, and compound E123 enhancer in CNCC. SV40 enhancer serves as positive control. Statistical comparisons to Empty control: SV40enh (p < 2.2×10^-16^), CTCF site (p = 0.64), E1 (p < 2.2×10^-16^), E2 (p < 2.2×10^-16^), E3 (p = 3.0×10^-4^), E123 (p < 2.2×10^-16^); n=8 independent transfections from 2 independent differentiations. Dunnett’s multiple comparisons test. **(C)** Schematic of TWIST1 degron-mediated depletion in CNCC for ChIP-seq (H3K27ac) and ATAC-seq at 0h (control) and 24h (depleted). **(D)** Genome browser tracks showing TWIST1 binding, H3K27ac, and accessibility at *TCOF1* enhancers E1-E3 in control and TWIST1-depleted CNCC. TWIST1 loss reduces both H3K27ac and accessibility at all three enhancers. **(E)** Schematic of E123 homozygous deletion. **(F)** TCOF1 transcript levels across differentiation in wild-type and E123 knockout cells by RT-ddPCR. E123 deletion subtly reduces TCOF1 levels in tCNCC2 (by 11%), tCNCC3 (by 30%), and CNCC (by 10%), but not in NE (Day5 neuroepithelium) or tCNCC1. n=3 independent differentiations for NE, n=5 independent differentiations for tCNCC1-3, n=6 independent differentiations for CNCC. ANOVA test followed by Dunn’s post-hoc test with Bonferroni adjustment.

To determine whether the identified enhancers regulate endogenous TCOF1 expression, we deleted all three enhancers homozygously (E123KO) in ESC and quantified *TCOF1* transcript levels across CNCC differentiation using RT-ddPCR (Fig. 3E,F). E123KO did not decrease *TCOF1* expression level in the NE and tCNCC1, which do not express TWIST1. In contrast, enhancer deletion led to a subtle, but significant reduction of *TCOF1* expression in the tCNCC2 (11% reduction), tCNCC3 (30% reduction) and passaged CNCC (10% reduction), which are all TWIST1-positive (Fig. 3F). Together, these data suggest that although *TCOF1* transcription does not strictly rely on enhancers – as may be expected from the housekeeping gene with a strong promoter – developmental enhancers nonetheless support *TCOF1* expression during CNCC mesenchymal transition.

### *TCOF1* enhancers support CNCC developmental transition

We next asked if enhancer deletion has functional consequences on CNCC differentiation in our *in vitro* model. We differentiated two distinct clonal E123KO ESC lines and collected cells at day 8 for scRNA-analysis. We observed selective loss of transitional populations (tCNCC 2/3), while pre-EMT NC remained intact (Fig. 4A,B, Fig. S4A,B). The strongest effect was observed for tCNCC3, consistent with stage-selective activity of the identified enhancers and largest effect on *TCOF1* expression. Pseudotime trajectory analysis confirmed developmental defect at the mesenchymal transition, similarly to what we have observed for TCOF1 dose perturbation (Fig. 4C). Moreover, in agreement with the scRNA-seq data and TCOF1 dosage perturbation data, we also observed impaired migration capacity in E123KO cultures (Fig. 4D,E).

**Figure 4.**
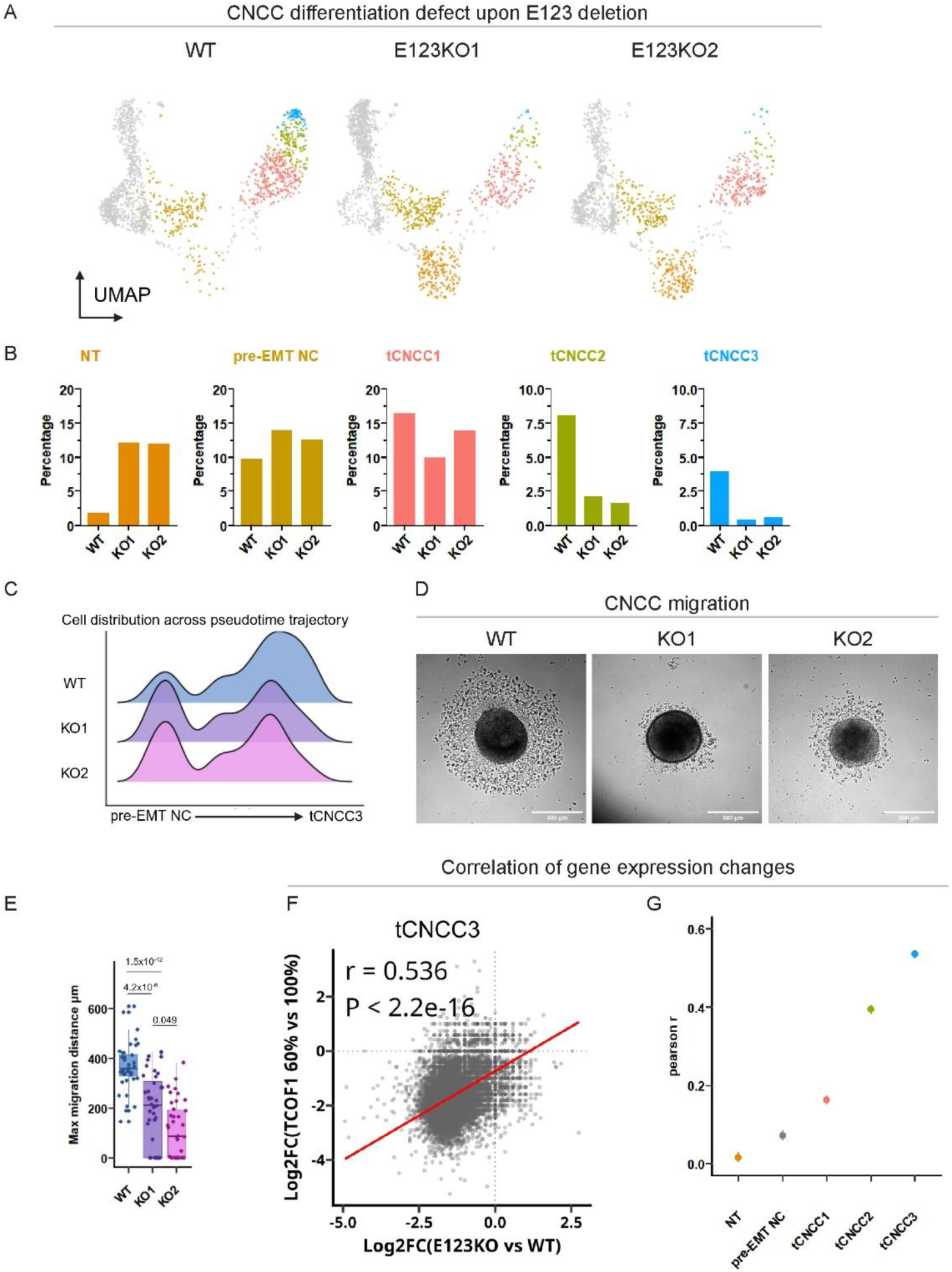
TCOF1 enhancers are essential for mesenchymal CNCC transition. **(A)** UMAP embedding comparing wild-type and two independent E123 knockout clones at day 8 shows selective loss of tCNCC2 and tCNCC3. **(B)** Quantification of cell type proportions shows E123 knockout reduces tCNCC2 and tCNCC3 while NT and pre-EMT NC remain intact and increase their relative proportion in the cell population. Data from one representative replicate (n=2 independent differentiations for each independent clonal line). **(C)** Distribution of cell populations across the pseudotime trajectory shows E123 knockout cells fail to progress from pre-EMT NC to later tCNCC stages. **(D)** Representative images of WT and E123 knockout (two independent clonal lines) in day 7 cultures. E123 knockout shows impaired cell migration compared to wild type. Scale bars, 500 μm. **(E)** Quantification of migration distance from neuroepithelial spheres. Boxplot from n=40 spheres across 3 independent differentiations. Error bars represent SEM. Kruskal-Wallis test followed by Dunn’s post-hoc test with Bonferroni adjustment. **(F)** Pearson correlation analysis comparing transcriptome-wide expression changes between E123KO vs. WT and TCOF1-depleted (60% protein) vs. control (100% protein) in tCNCC3. Each dot represents one gene (n=19278). Log₂ fold-changes were calculated using DESeq2 from 2 independent differentiations per condition. P-value from t-distribution (H₀: ρ=0, df=n-2). **(G)** Pearson correlation coefficients of transcriptome-wide expression changes from panel F and FigS4 across differentiation stages, showing increased correlation during tCNCC2 and tCNCC3 transition, correlating with peak enhancer activity. Error bars representing 95% confidence intervals (CI), calculated using Fisher’s z-transformation.

To further assess whether E123KO phenocopies reduced TCOF1 dosage at the molecular level, we compared transcriptional effects of E123KO versus TCOF1 depletion across the developmental trajectory, from NT to tCNCC3, by correlating gene expression changes between conditions in each population. E123KO showed significant correlation with TCOF1 depletion in tCNCC2 and tCNCC3, but minimal correlation in NT, pre-EMT NC or tCNCC1 stages, where enhancers show low/no activity and their deletion has minor effects on transcriptome in general (Fig. 4F, Fig. S4C). The correlation coefficient between transcriptomic effects of two perturbations steadily increased with the developmental progression, along with enhancer opening (Fig. 4G). Altogether, this stage-specific transcriptional and phenotypic convergence between TCOF1 dosage reduction and E123KO, indicates that *TCOF1* regulation by enhancers is crucial for proper CNCC development.

### Enhancers buffer *TCOF1* expression against transcription factor fluctuations

Although the yield of tCNCC3 is severely affected in E123KO cells, we were able to isolate ‘escapee’ tCNCC3 cells and culture them in the passage media we typically use for long-term maintenance of CNCC. The wild-type passaged CNCC have strong mesenchymal characteristics and high level of TWIST1 expression, and consistent with these features, the E1-E3 enhancers are accessible and active in these cells (Fig. 3A,B). We were therefore surprised that in passaged CNCC with the E123KO, TCOF1 expression reduction was only ∼10%, compared to the 30% reduction seen in tCNCC3 (Fig. 3F). Notably, CNCC are passaged under culture conditions that stabilize the mesenchymal state and allow for its long-term maintenance (*28*), whereas tCNCC undergo dynamic fate changes, actively delaminate and migrate away from the neuroepithelial spheres. We hypothesized that the stronger requirement for E1-E3 in tCNCC3 may be caused by less stable regulatory inputs into the *TCOF1* promoter during these dynamic developmental transitions, and that the key role of the enhancers may be to buffer expression against fluctuations in promoter-regulating transcription factors.

To explore this possibility, we focused on MYC, which directly binds the *TCOF1* promoter (*39*) (Fig. S5). MYC is a master regulator of ribosome biogenesis and other housekeeping genes (*7, 40*), and plays an essential role in neural crest development (*41, 42*). MYC is also a direct target of WNT signaling, a key developmental signal provided by the dorsal neural tube that regulates neural crest development (*18, 43*). More broadly, MYC protein levels are known to be inherently variable, responding dynamically to developmental signals, metabolic status, and environmental cues; such variable expression can be the basis of cell competition observed in many developmental systems (*44–47*) (Fig. 5A). We reasoned that if enhancers buffer against variability in levels of promoter-regulating transcription factors, TCOF1 expression should remain stable across MYC fluctuations in wild-type cells but become sensitized to MYC changes in E123KO cells. To mimic MYC fluctuations in passaged CNCC, which are grown under stable conditions and show minimal dependency on enhancers for TCOF1 expression, we employed a MYC-targeting molecular glue degrader (*48*). With this degrader, we achieved a dose-dependent MYC depletion, validated by intracellular flow cytometry (Fig. 5B). We then assayed endogenous TCOF1 transcript levels by RT-ddPCR across MYC titrations. This analysis revealed striking differences between genotypes, where wild-type CNCC exhibited robust buffering and TCOF1 expression remained stable across moderate MYC depletion, declining sharply only upon severe MYC loss (Fig. 5C,E). In contrast, E123KO CNCC lost this buffering capacity, showing immediate, high sensitivity to MYC reduction (Fig. 5D,E).

**Figure 5.**
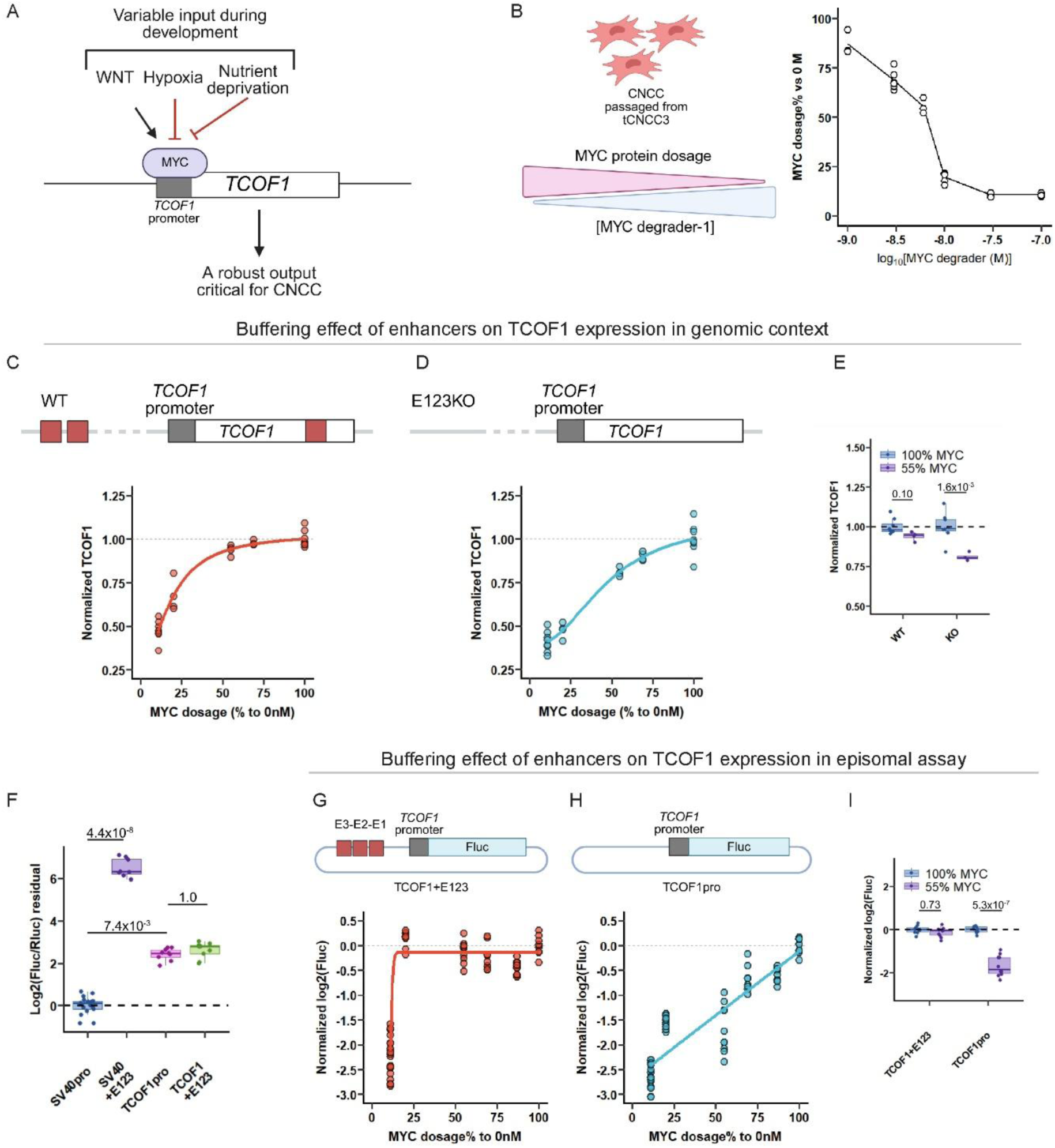
TCOF1 enhancers buffer against fluctuations in MYC dosage. **(A)** Schematic showing MYC regulation by developmental (e.g. WNT), environmental and metabolic signals and MYC binding to the TCOF1 promoter. **(B)** Left, schematic of MYC molecular glue degrader treatment. Right, flow cytometry quantification of dose-dependent MYC depletion in CNCC from 3 independent differentiations. **(C)** Endogenous locus buffering assay: *TCOF1* transcript levels in response to MYC titration in wild-type CNCC show buffered response until severe depletion. n=4 independent differentiations. Fitted non-linear LL4 model. **(D)** Endogenous locus buffering assay: *TCOF1* transcript levels in response to MYC titration in E123 knockout CNCC show unbuffered sensitivity. n=4 independent differentiations. Fitted non-linear LL4 model. **(E)** *TCOF1* expression at 55% MYC in wild-type vs. E123 knockout CNCC. E123 knockout shows significant reduction while wild-type maintains robust expression. n=4 independent differentiations. Dunnett’s multiple comparisons test. **(F)** Dual luciferase assay comparing E123 effects on SV40 minimal promoter vs. *TCOF1* promoter activity, demonstrating lack of signal amplification by coupling enhancers to the *TCOF1* promoter. Two assay sets were performed with a shared SV40 promoter control for cross-assay normalization: SV40 promoter vs. SV40+E123 (n=8 independent transfections across 2 independent differentiations); *TCOF1* promoter vs. TCOF1+E123 (n=9 independent transfections across 3 independent differentiations). Statistical comparisons: SV40pro vs. SV40+E123 (p = 4.4×10⁻⁸), SV40pro vs. TCOF1pro (p = 7.4×10⁻³), TCOF1pro vs. TCOF1+E123 (p = 1.0); ANOVA test followed by Dunn’s post-hoc test with Bonferroni adjustment on shared-control anchored data. **(G)** Episomal luciferase reporter assay: reporter activity driven by *TCOF1* promoter + enhancer (TCOF1+E123) in response to MYC titration shows buffered, threshold-protected output. n=10 independent transfections across 2 independent differentiations. Fitted non-linear LL4 model. **(H)** Episomal luciferase reporter assay: reporter activity driven by *TCOF1* promoter alone (TCOF1pro) in response to MYC titration shows linear, unbuffered sensitivity. n=10 independent transfections across 2 independent differentiations. Fitted linear model. **(I)** Firefly luciferase at 55% MYC compared to TCOF1+E123 vs. TCOF1pro demonstrates enhancer-mediated buffering. n=10 independent transfections across 2 independent differentiations. Dunnett’s T3 test.

To validate these findings with an orthogonal approach, we constructed luciferase reporters containing either the *TCOF1* promoter alone (TCOF1pro) or coupled with compound E123 enhancer elements (TCOF1+E123). Interestingly, while SV40 minimal promoter was strongly (∼90 fold) boosted by the presence of E123, the *TCOF1* promoter showed no significant change in activity (Fig. 5F). Despite comparable activity in the unperturbed state, MYC titrations revealed a dramatic difference in the dose response curve shape between the two reporters: TCOF1+E123 maintained robust, buffered expression until MYC dose was reduced to 25% of the endogenous levels, while TCOF1 promoter alone exhibited linear, unbuffered sensitivity (Fig. 5G-I). These experiments establish that enhancers buffer TCOF1 expression by converting a sensitive, dose-dependent promoter regulation by MYC into a buffered, threshold-governed output. This buffering explains the differential enhancer dependence observed across developmental stages: E123 becomes critical during tCNCC3 transition when robust TCOF1 expression must be maintained against variable regulatory inputs but becomes less essential under optimal culture conditions such as those in passaged CNCC.

### Enhancer-mediated buffering represents a generalizable regulatory mechanism for dosage-sensitive housekeeping genes

To test whether the enhancer-mediated buffering mechanism uncovered for *TCOF1* generalizes to other dosage-sensitive housekeeping genes, we developed a systematic approach to identify candidate genes with *TCOF1*-like properties. We integrated two genome-wide metrics: (i) pHaplo scores—quantifying haploinsufficiency sensitivity from human genetic scans (*9*)—and (ii) expression coefficient of variation across our developmental stages (ESC, NE, tCNCC1+2, and passaged CNCC) to identify genes whose expression is relatively stable across differentiation (Data S1). Applying both criteria (pHaplo > 0.8, CV < 0.06) revealed *TCOF1* as a typical example, consistent with a dosage-sensitive housekeeping gene. Using these thresholds, we selected 13 additional promoters with similar properties, together with *TCOF1* forming our dosage-sensitive housekeeping (DSH) gene set (n=14, Fig. 6A). Consistent with their high pHaplo scores, these DSH genes are linked to a variety of human disorders spanning multiple organ systems (Data S2), illustrating that genes can simultaneously exhibit housekeeping expression patterns— characterized by broad tissue distribution and low expression variance—while maintaining dosage sensitivity.

**Figure 6.**
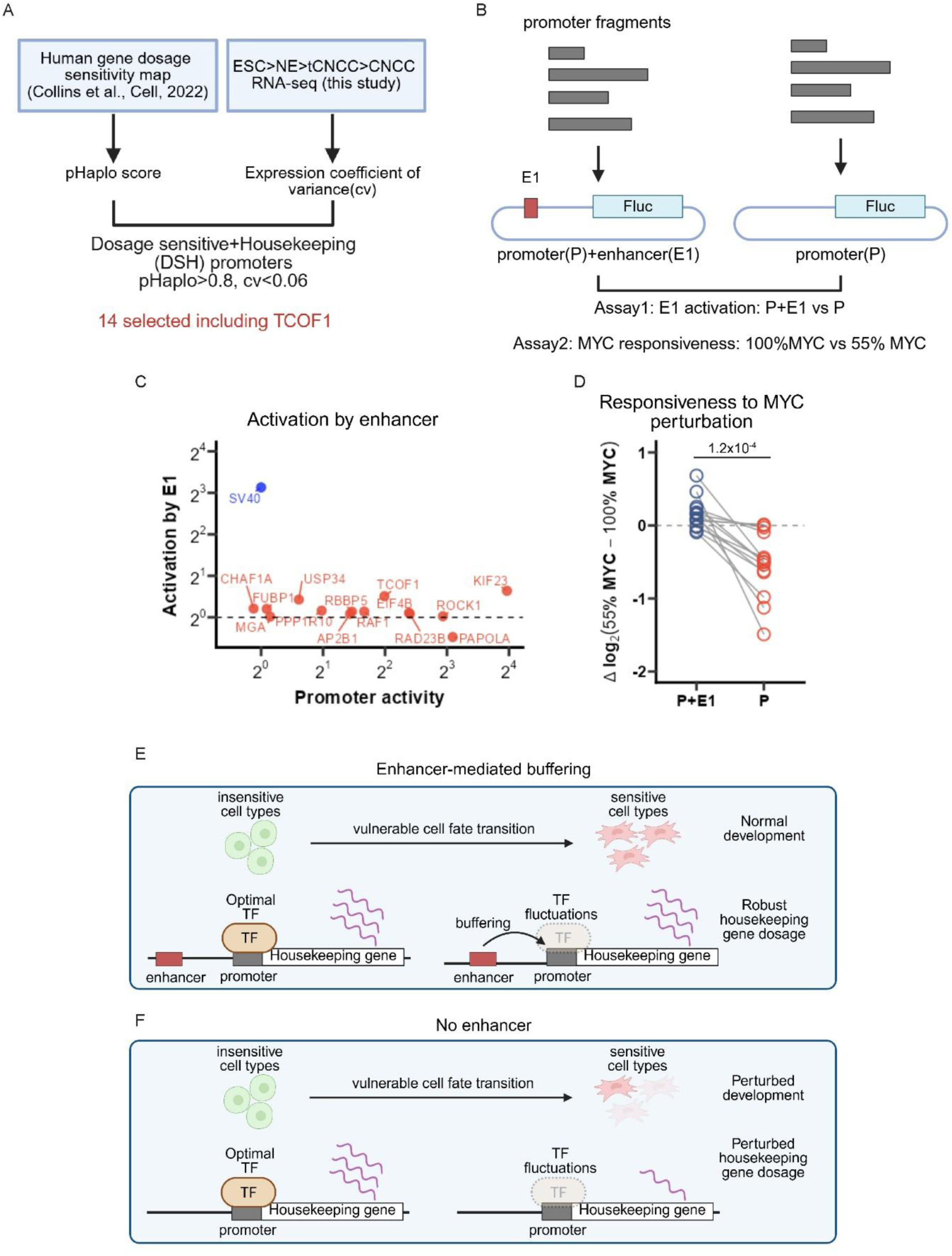
Enhancer buffering generalizes across many dosage-sensitive housekeeping genes. **(A)** Selection criteria for dosage sensitive housekeeping (DSH) genes, using pHaplo scores > 0.8 and expression coefficient of variance < 0.06, as measures of dosage sensitivity and broad expression across cell types, respectively. **(B)** Schematic of luciferase reporter assay: promoter-only(P) vs. promoter+TCOF1-E1(P+E1). **(C)** Dual luciferase assay comparing promoter activity alone vs. promoter+E1 for all 14 DSH promoters and control SV40 promoter in CNCC. DSH promoters show minimal amplification; SV40 promoter shows strong amplification; result integrated from 2 independent differentiations. **(D)** Promoter responsiveness to MYC depletion (expression at 55% MYC/100% MYC) comparing promoter-only vs. promoter+E1 constructs. Dosage-sensitive housekeeping (DSH) promoters alone are MYC-sensitive; P+E1 coupling provides buffering. n=14 DSH promoters with 3 independent differentiations. Statistical significance assessed by paired Wilcoxon signed-rank test. **(E)** Schematics of enhancer buffering function at dosage-sensitive housekeeping genes **(F)** Loss of enhancers at dosage-sensitive housekeeping genes manifests during vulnerable cell fate transitions associated with fluctuations in promoter-regulating TFs.

We first tested whether DSH promoters respond to enhancers through classical transcriptional amplification. Each of the 14 DSH promoters and the SV40 control promoter were cloned into luciferase reporters either alone or coupled to the *TCOF1*-E1 enhancer, then assayed in CNCC (Fig. 6B). DSH promoters showed minimal amplification when coupled to E1, similar to previously reported work (*3, 49*). Importantly, this lack of amplification was independent of a baseline promoter strength, suggesting that regardless of their activity, these promoters operate at or near saturation. In contrast, the SV40 minimal promoter showed robust amplification with E1, consistent with canonical enhancer function (Fig. 6C).

To test whether enhancers provide buffering functions to DSH promoters, we repeated the reporter assay under MYC level reduction to 55% of endogenous levels. In these conditions, DSH promoters alone showed significant expression loss, but coupling these same promoters to E1 maintained near-complete stabilization of expression, demonstrating robust buffering (Fig. 6D). Thus, enhancer-mediated buffering of housekeeping gene expression represents a generalizable regulatory strategy distinct from the canonical transcriptional amplification by enhancers.

## Discussion

Our findings have important implications for understanding the role of distal enhancers in the regulation of housekeeping genes. Minimal amplification of the transcriptional output from the housekeeping promoters in the presence of enhancers has been previously observed in the massively parallel reporter assays, and sometimes interpreted as an inherent lack of regulatory compatibility of these promoters with developmental enhancers (*3, 4*). However, our results suggest that under optimal conditions, the housekeeping promoters operate at or near activity saturation, and thus their expression is only weakly responsive to boosting by enhancers. Interestingly, this lack of enhancer-mediated boostability holds for promoters across a wide range of inherent strengths. However, fluctuations in the promoter-regulating transcription factors such as MYC unmask the requirement for enhancer-mediated regulation. Thus, rather than strictly amplifying expression, enhancers reshape transcription factor dosage responses at housekeeping promoters, buffering against regulatory variability (Fig. 6E,F). We postulate that such mechanism may be particularly biologically relevant for dosage-sensitive housekeeping genes during vulnerable cell fate transitions characterized by dynamic fluctuations of promoter-regulating transcription factors. Such factors, including MYC, SP1, NRF1 or GABPA (NRF2) are broadly expressed across cell types, but nonetheless sensitive to growth and metabolic cues (*43, 46, 47, 50, 51*). Our findings exemplify this principle: while broadly expressed, *TCOF1* requires CNCC-specific enhancers that convert its promoter’s linear, sensitive response to MYC into a buffered, non-linear output, in turn maintaining robust expression during mesenchymal fate specification (Fig. 6E,F).

The biological necessity of this buffering stems from TCOF1’s extreme dosage sensitivity in CNCC—even modest protein level reduction of ∼20% impairs rRNA synthesis and mesenchymal fate specification. This hypersensitivity contrasts sharply with POLR1 subunits, which exhibit robust buffering capacity against dosage reduction beyond a two-fold change in levels. This differential sensitivity resolves the longstanding genetic mystery of TCS: *TCOF1* mutations, including complete loss-of-function alleles, account for vast majority of cases because TCOF1 is rate-limiting for rRNA synthesis. In contrast, combined *POLR1A/B/C/D* mutations account for ∼10% of cases and are predominantly missense variants likely acting through dominant-negative mechanisms rather than simple haploinsufficiency (*15, 22, 23, 52*). TCOF1’s dosage sensitivity further compounds with selective vulnerability of CNCC to perturbations in rRNA synthesis, together providing a mechanistic explanation for specificity of TCS craniofacial pathology.

TCOF1’s extreme dosage sensitivity may also help explain substantial interfamilial and intrafamilial phenotypic variability that characterizes TCS patients (*16, 53*). While heterozygous *TCOF1* loss is inherently pathogenic, phenotypic severity may be exacerbated or alleviated depending on the buffering capacity of the unaffected allele and genetic/environmental modifiers. Individuals with stronger enhancer activity at the unaffected *TCOF1* locus, favorable genetic variants in MYC, WNT, or p53 pathways, or stable maternal metabolic conditions may maintain *TCOF1* expression at higher levels within the pathogenic range, resulting in milder phenotypes, whereas those with compromised buffering face more severe manifestations. Our findings suggest that prenatal interventions stabilizing MYC signaling during critical developmental windows could mitigate TCS severity, and that this principle could potentially apply to other housekeeping genes associated with developmental haploinsufficiencies. Such interventions could include maternal dietary supplementation linked to pathways known to stabilize MYC, including amino-acid sensing, acetate metabolism and direct PI3K/mTORC1 activation (*50, 54, 55*).

The enhancer-mediated buffering at housekeeping promoters draws an interesting parallel to the phenomenon of redundancy between ‘shadow’ and primary enhancers described at developmental genes in *Drosophila*. Shadow enhancers are secondary enhancers that share the same spatiotemporal activity and gene target with the primary enhancers and appear functionally redundant with them under normal conditions (*56–58*). Yet, the requirement for shadow enhancers is unmasked under environmental stress, where they confer robustness to developmental phenotypic outputs (*59, 60*). In both mechanisms – enhancer redundancy and buffering of housekeeping promoters – the buffering function of a regulatory element is cryptic under optimal conditions and only revealed when the system is challenged. Interestingly, however, the precise cis-regulatory architecture through which robustness is achieved is distinct in the two mechanisms. At developmental genes, robustness emerges between multiple enhancers: when one enhancer is compromised, another enhancer (with similar spatiotemporal activity but distinct cis-regulatory wiring) compensates to sustain transcriptional output (*60–62*). At dosage-sensitive housekeeping genes, as we show here, robustness instead emerges from the interaction between a strong, saturable promoter and its enhancers, which together produce non-linear, buffered transcriptional response to fluctuations in promoter-regulating transcription factors. Together, these observations suggest that enhancer-mediated buffering—whether achieved through enhancer redundancy or through TF dose-response reshaping at promoters—may represent a widespread and fundamental regulatory principle ensuring robustness of gene regulation.

## Acknowledgments

We thank Wysocka lab members, as well as Lei Li, for stimulating discussions and critical input into the manuscript. Flow cytometry analysis and sorting were done with Stanford University Stem Cell Institute flow cytometry core. Illustrations in Figures 1A, 2A, 2J, 3A, 3C, 3E, 5A, 5B, 5C, 5D, 5G, 5H, 6A, 6B, 6E, 6F, S1D, S3A are created partially or fully with BioRender.com.

## Funding

Damon Runyon Cancer Research Foundation Fellowship DRG 2505-23 (L.I.)

Howard Hughes Medical Institute (J.W.)

Nomis Foundation (J.W.)

National Institutes of Health grant R35 GM131757 (J.W.)

California Institute for Regenerative Medicine grant DISC0-17487 (J.W.)

## Author contributions

Conceptualization: CN, JW

Methodology: CN, LI, TS, JW

Investigation: CN, LI, TS, JW

Visualization: CN, JW

Funding acquisition: JW

Project administration: CN, JW

Supervision: JW

Writing – original draft: CN, JW

Writing – review & editing: CN, LI, TS, JW

## Competing interests

The authors declare no competing interests.

## Data, code, and materials availability

All data needed to evaluate the conclusions in this study are present in the main text, the supplementary materials, and/or the Gene Expression Omnibus

Accession numbers of reanalyzed publicly available datasets are listed in Table S2.

## Supplementary Materials

Materials and Methods

Figs. S1 to S5

Tables S1 to S2

Data S1 to S2

## Materials and Methods

### Stem cell culture

H9 human embryonic stem cells were obtained from WiCell. hESC were cultured in mTESR-plus medium (Stem Cell Technologies, 100-0276) on matrigel-coated plates (Corning, 354277). Cells were passaged every 3-5 days after dissociation with ReLeSR (Stem Cell Technologies, 100-0483). Routine mycoplasma tests were performed with Universal Mycoplasma Detection Kit (ATCC, 30-1012K).

### CNCC induction and passaging

To model CNCC induction, in Aggrewell 800 plates (Stem Cell Technologies, 34815), 3×10^5 ESC in 2 mL CNCC induction medium(1:1 ratio of DMEM-F12 and Neurobasal, 0.5× Gem21 NeuroPlex supplement with vitamin A, Gemini, 400-160, 0.5× N2 NeuroPlex supplement, Gemini, 400-163, 1× antibiotic–antimycotic, 0.5× Glutamax, 20 ng/ml bFGF, PeproTech, 100-18B, 20 ng/ml EGF, PeproTech, AF-100-15 and 5 μg/ml bovine insulin, Gemini Bio-Products, 700-112P) were added per well to achieve 1000 cells per neuroectoderm(NE) sphere. During the formation period, the medium was changed 75% every day with an electronic auto dispenser at slowest pipetting speed. On day 4 of induction, NE spheres were collected and transferred into 6-well plates containing 2 mL of fresh CNCC Induction Medium per well. By day 6, the spheres attached to the culture surface, and migratory CNCC began to emerge from the central NE core. This culture, which captures the delamination process, is referred to as the NE-CNCC transitioning model. The medium was replaced on day 6. For assays studying this transitional state, cultures were harvested on day 8. To generate a purified population of mesenchymal CNCC, the NE-CNCC transitioning models were maintained until day 11, with daily medium changes performed from day 8 onward. On day 11, migrated CNCC were passaged onto fibronectin-coated 6-well plates at 1×10^6/well, with CNCC maintenance medium (1:1 ratio of DMEM-F12 and Neurobasal, 0.5× Gem21 NeuroPlex supplement with vitamin A, Gemini, 400-160, 0.5× N2 NeuroPlex supplement, Gemini, 400-163, 1× antibiotic–antimycotic, 0.5× Glutamax, 20 ng/ml bFGF, PeproTech, 100-18B, 20 ng/ml EGF, PeproTech, AF-100-15, 1 mg/ml BSA). The passaged CNCC were counted as P1. On day 13, P1 CNCC were counted and passaged onto fibronectin-coated 6-well plates at 8×10^5/well, with CNCC-BC medium(CNCC maintenance medium + 1ng/ml BMP2, PeproTech, 120-02 + 3 μM CHIR-99021, Selleck Chemicals, S2924). After transition to CNCC-BC medium, CNCC at subsequent passages were plated at 8×10^5/well per well of a 6-well plate. CNCC were then passaged twice to P4CNCC.

### Dissociation of differentiating CNCC and scRNA-seq

NE-CNCC transitioning models from 1 well in 6-well plate(∼100x spheres plated on day 4), including spheres and migrated cells, were dissociated in ACCUTASE (Stem Cell Technologies, 07920) for 10min at 37 °C. scRNA-seq libraries were generated using the Illumina Single Cell 3’ RNA Prep T2(Illumina, 20135689). For each sample, an estimated 2000 cells were captured. Libraries were sequenced with Novaseq X platform.

### scRNA-seq data processing and analysis

Reads from scRNA-seq were aligned to the GRCh38 human reference genome and the cell-by-gene count matrices were produced using illumina’s proprietary software DRAGEN with default parameters. Data were analyzed using the Seurat R package v4 using R v.4.6. Data from all 12 individual samples were merged. Cells were filtered on the basis of unique molecular identifier (UMI) counts (>500), the number of detected genes (>500), the log10 genes per UMI (>0.8), and the fraction of mitochondrial genes (<0.2). Cell-cycle related genes were excluded from the set based on AnnotationHub. Seurat project was then split, and individual samples were normalized, scaled and centered by SCTransform(), with percent.mt and cell cycle regressed. Principal Component Analysis (PCA) was performed using the Seurat function RunPCA (). The first 30 PCs were used to integrate the different treatment conditions and genotypes in the dataset using Harmony with max iteration as 50. We performed UMAP using RunUMAP () with reduction method as “harmony” and otherwise default settings. Cells were clustered in PCA space using FindNeighbors (), followed by FindClusters with resolution = 1.0. Differentially regulated genes in each cluster were identified using FindAllMarkers () with logFC threshold at 0.25. The top significant differentially regulated genes, along with curation by pre-knowledge, were selected for dot plot and heatmap visualization. We subset cells belonging to the pre-EMT, tCNCC1, tCNCC2 or tCNCC3, and performed differential gene expression analysis using DESeq2 on the sample-level aggregated counts (pseudobulk), to compare WT vs KO and control vs moderate TCOF1 depletion. All detected genes were then used for correlation analysis. Pseudotime trajectory analysis was performed using Monocle3 v0.2.0 with default parameters. The root of the trajectory was defined as the endpoint exhibiting the highest WNT1 expression. Prior to trajectory inference, cells were downsampled to ensure equal representation across all experimental conditions.

### Live cell flow cytometry analysis and sorting

NE-CNCC transitioning models from 1 well in 6-well plate(∼100x spheres plated on day 4), including spheres and migrated cells, were dissociated in ACCUTASE (Stem Cell Technologies, 07920) for 10min at 37 °C. Dissociated cells were pelleted at 300g x 3min at 4 °C, resuspended with DPBS-3%KSR, and counted using a Countess Automated Cell Counter (Invitrogen). For analysis purposes, cells were stained with CD49D-APC(Biolegend 304308) or CD49D-PE-CY7(Biolegend 304314), CD10-BV421(BD 659449) and/or ERBB3-488(R&D Systems FAB3481G) with 1:100 in DPBS-3%KSR for 30min at 4 °C. Cells were then washed once with DPBS-3%KSR and resuspended in DPBS-3%KSR at 5×10^6/mL for flow cytometry analysis(BD FACSymphony A5, BD LSRFortessa) and sorting(BD FACSMelody).

## Intracellular flow cytometry analysis

NE-CNCC transitioning models were dissociated, counted and resuspended in DPBS-3% KSR, with necessary surface staining of CD49D-APC and CD10-BV421 as mentioned in the section of live cell flow cytometry analysis and sorting. After staining, cells were washed twice with DPBS-3% KSR, and fixed 4% PFA at RT for 10min. Cells were then washed twice with DPBS-3% BSA and permeabilized with DPBS-0.5%BSA-0.1% Triton X-100 at RT for 15min. Cells were then incubated with primary antibodies in DPBS-0.5% BSA for 16-24h at 4 °C. Primary antibodies used in this study include anti-TCOF1 (Proteintech 11003-1-AP), anti-ERBB3-488(R&D Systems FAB3481G), anti-V5-488(Thermo 740058MP488).

After primary antibody, cells were washed once with DPBS-0.5% BSA before incubating with secondary antibodies in DPBS-0.5% BSA for 1h at 4 °C. Cells were then washed twice and resuspended with DPBS-0.5% BSA for flow cytometry analysis (BD FACSymphony A5, BD LSRFortessa).

### cDNA preparation and reverse transcription digital droplet PCR (RT-ddPCR)

Total RNA was extracted from ESC and passaged CNCC for wild-type or enhancer mutant cells using Trizol reagent (Invitrogen) on plate, followed by Direct-zol RNA microprep kit (Zymo R2060) with on-column DNase I digestion. For tCNCC1(CD49D+/ERBB3+), tCNCC2(CD49D+/ERBB3-/CD10-), tCNCC3(CD10+), 1×10^5 cells were sorted from NE-CNCC transitioning models and total RNA was extracted with Trizol-LS reagent (Invitrogen), followed by Quick-RNA microprep kit (Zymo) with on-column DNase I digestion. 100-1000ng RNA was used to generate cDNA using the SuperScript Vilo IV MasterMix (Invitrogen). ddPCR reactions were performed using diluted cDNA (equivalent to 4ng of total RNA), 500nM primers and 250nM probes and 1X ddPCR Supermix for probes (no dUTP, BioRad). ddPCR droplets were generated using the QX200 Droplet Generator (BioRad) and droplets were read using QX200 Droplet Reader (BioRad) and analyzed using the QuantaSoft Software (BioRad). Primers and probes are in Table S1.

### ATAC-seq sample collection and library preparation

6×10^4 cells from NE(CD49D-/CD10-), tCNCC1(CD49D+/ERBB3+), tCNCC2(CD49D+/ERBB3-/CD10-), tCNCC3(CD10+) were sorted from NE-CNCC transitioning models, pelleted at 500xg for 5 min at 4 °C and resuspended in ATAC-resuspension buffer (10 mM Tris-HCl pH 7.4, 10 mM NaCl, 3 mM MgCl2 in sterile water) containing 0.1% NP-40, 0.1% Tween20 and 0.01% digitonin and incubated on ice for 3 min. Following wash-out with cold ATAC-resuspension buffer containing 0.1% Tween20, cells were pelleted and resuspended in 50 μl transposition mix (25 μl 2× TD buffer, 2.5 μl transposase (100 nM final), 16.5 μl PBS, 0.5 μl 1% digitonin, 0.5 μl 10% Tween20, 5 μl H2O) and incubated for 30 min at 37 °C with shaking. The reaction was purified using the Zymo DNA Clean & Concentrator kit and PCR-amplified with NEBNext High-Fidelity 2× PCR Master Mix (NEB, M0541L). Libraries were purified by two rounds of double-sided size selection with AMPure XP beads (Beckman Coulter, A63881), with the initial round of 0.5× sample volume of beads, taking the supernatant, followed by a second round with 1.3× initial volume of beads. Pooled libraries were sequenced using the Novaseq X platform.

### ATAC-seq data processing

Paired-end reads were aligned to the hg38 reference genome using BWA 0.7.19-r1273 with default parameters. Aligned reads were sorted and compressed to BAM format using samtools 1.22.1. BAM files were indexed with samtools index for downstream analysis. For visualization, genome-wide coverage tracks were generated and normalized to library size as follows. Coverage was calculated using bedtools v2.31.1 with RPKM (Reads Per Kilobase per Million mapped reads) normalization, where a scaling factor of 1,000,000 divided by the total number of mapped reads was applied. The resulting bedGraph files were sorted and converted to bigWig format using UCSC bedGraphToBigWig for visualization in genome browsers.

### Bulk RNA-seq sample collection, sequencing and analysis

Cells were harvested by TRIzol (Invitrogen, 15596018) and stored at −80°C until processing. Total RNA was extracted by Direct-zol RNA microprep kit (Zymo R2060). RNA was sequenced with novogene EukmRNAseq-NovaSeqXPlus-6Gb. RNA-seq data were mapped using salmon quant and analyzed by DESeq2.

### CRISPR-mediated gene editing

For the enhancer knock-out, 8×10^5 ESC were nucleofected with two Cas9–sgRNA RNP complex(100pmol Sp-Cas9 HiFi, IDT, 300 pmol sgRNA, IDT) for upstream and downstream cut, using the Lonza 4D Nucleofection system. For homology-directed recombination (HDR) of FKBPV and FKBPV-mNG, 8×10^5 ESC were nucleofected with a Cas9–sgRNA RNP complex(100pmol Sp-Cas9 HiFi, IDT, 300 pmol sgRNA, IDT) and 1pmol of dsDNA donor, using the Lonza 4D Nucleofection system. Cells were plated on matrigel-coated 6-well plate with mTeSR Plus+CET (50nM Chroman 1, 5 µM Emricasan, 0.7 µM trans-ISRIB). 24h after nucleofection cells were changed with mTeSR Plus without CET. Cells were further maintained to day 4 and were sorted into matrigel-coated 96-well plate at 1 cell/well mTeSR Plus+CET. 12 days after plating, colonies grown in 96-well plate were expanded into 24-well plate. Colonies were screened by PCR and Sanger sequencing. Desired mutant and knock-in clones were expanded. Sequences of sgRNA and PCR primers are available in Table S1.

### Migratory distance analysis

Individual NEs on Day 6 were plated into 96-well flat-bottom plate at 1 sphere/well. 24h after plating, images of NE-CNCC transitioning models were taken with Nikon Eclipse Ti-E at x10 magnification objective. Distance values were obtained by quantifying individual NE-CNCC migration on ImageJ, which measured distance in µm.

### dTAGv-1-mediated protein degradation

For POLR1A degradation in ESC and passaged CNCC, cells were treated with DMSO vehicle or dTAGv-1 at specified concentration for 24 h prior to assay. For POLR1A, POLR1B and TCOF1 degradation during CNCC differentiation, dTAGv-1 was added to CNCC induction medium at specified concentrations from day 4 to day 8 of NE-CNCC transitioning models, with medium changes every 2 days maintaining dTAGv-1 concentration. For rRNA synthesis assay during CNCC differentiation, dTAGv-1 was added to CNCC induction medium at specified concentrations on day 8 of NE-CNCC transitioning models for 24h treatment. Treated cultures were then processed for flow cytometry analysis, scRNA-seq as described above.

### 5-EU incorporation assay for rRNA synthesis

At the time of the assay, cells were incubated with complete medium(mTESR-PLUS for ESC, CNCC induction medium for differentiating model, CNCC-BC for P4CNCC) with 0.5 mM EU(clickchemistrytools) for 1 h. Cells were then washed twice with DPBS, dissociated with Accutase at 37°C for 10min, washed once with DPBS-3% KSR, counted using a Countess Automated Cell Counter. 1×10^6 cells per assay were fixed with 4% formaldehyde (v/v) in DPBS for 15 min and permeabilized with 0.1% saponin in DPBS for 15 min. After permeabilization, cells were washed once with 3%BSA in DPBS, and then incubated with click-reaction cocktail made with Click-&-Go Cell Reaction Buffer Kit and 2.5μM AFDye 568 Azide (Click Chemistry Tools) for 30 min. After reaction, cells were washed once with 3%BSA in DPBS, and resuspended with DPBS-0.5% BSA for flow cytometry analysis (BD FACSymphony A5, BD LSRFortessa).

### MYC degradation

For MYC degradation experiments, P4CNCC were treated with MYC degrader 1 (MCE, HY-162318) at indicated concentrations in CNCC-BC medium for 20 h prior to flow cytometry analysis, RNA extraction and luciferase assay.

### Luciferase assay

CNCC were transfected with 0.1 ng pRL renilla control plasmid, 2 ng modified pGL3 reporter plasmid, and 17.9 ng carrier plasmid (pUC19) per well of a 96-well plate. For enhancer activation assay, cells were lysed 24 h after transfection and assayed with the Dual-Luciferase Reporter assay kit (Promega, E1960). For MYC responsiveness assay, cells were lysed 20 h after MYC degrader 1 treatment and assayed with the Luciferase Assay System Kit (Promega, E1500).

---

**Figure S1.**
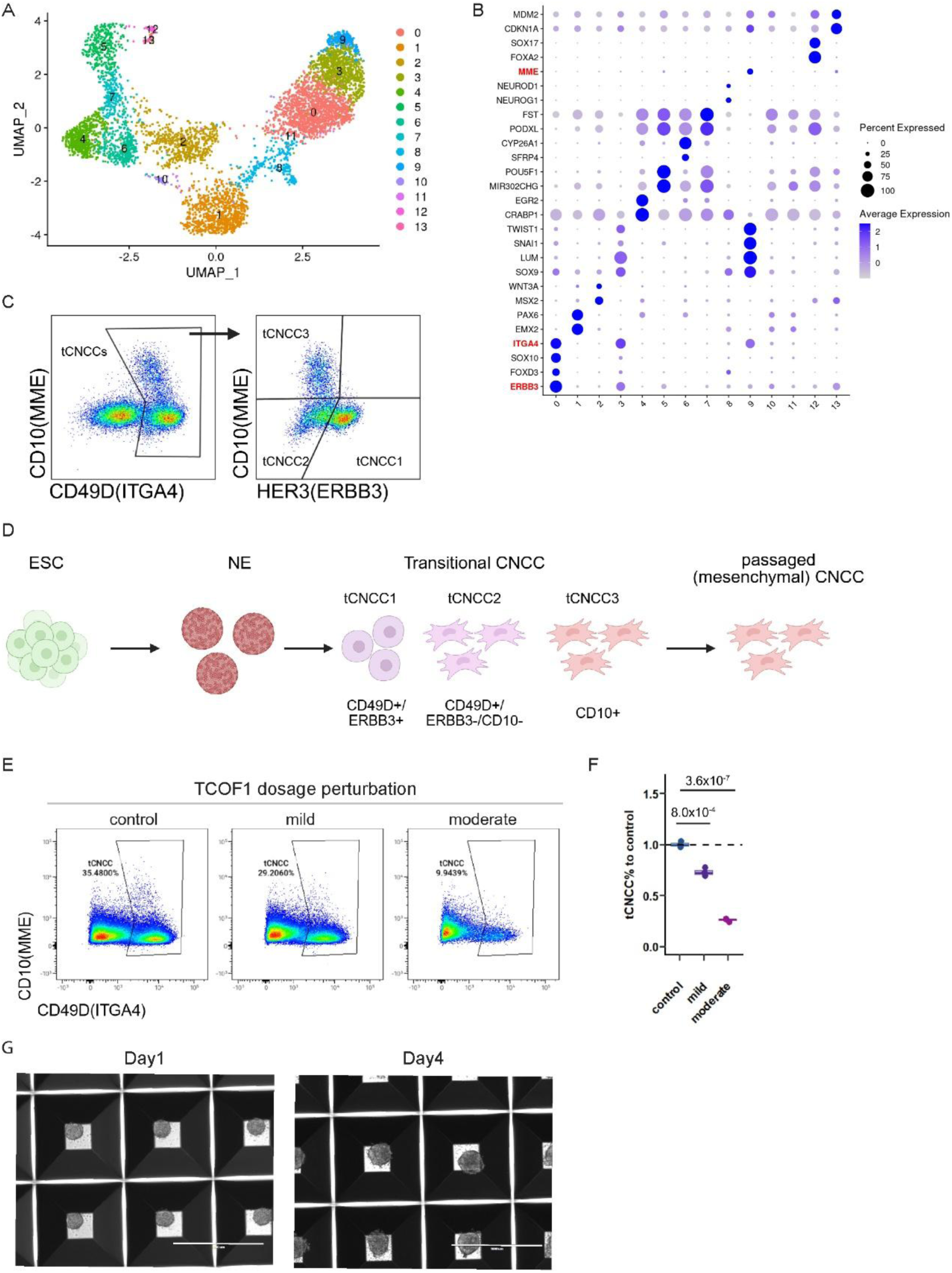
Comprehensive single-cell analysis of day 8 CNCC differentiation and TCOF1 dosage response. **(A)** Complete UMAP embedding of day 8 cells reveals discrete populations spanning neural tube (cluster 1), pre-EMT neural crest (cluster 2), three sequential transitional CNCC states (tCNCC1-3, clusters 0, 3, 9), pluripotent/early neuroectoderm (clusters 5, 7, 4, 6), neuronal cells (cluster 8), unknown or stressed cells (clusters 10, 11, 12, 13). **(B)** Comprehensive marker gene expression panel used to define cluster identities across the NT-to-CNCC developmental trajectory. Surface markers used for flow cytometry isolation in this study (ERBB3, ITGA4, MME) are marked in red. **(C)** Representative flow cytometry gating strategy for prospective isolation of tCNCC1, tCNCC2, and tCNCC3 populations using surface marker combinations identified in this work. **(D)** Schematic overview of cell types and developmental stages analyzed throughout this study. **(E)** Representative flow cytometry plots showing tCNCC population percentages (combined tCNCC1-3) in control (100% TCOF1), mild TCOF1 depletion (82% TCOF1), and moderate TCOF1 depletion (60% TCOF1) conditions. **(F)** Quantification of tCNCC percentage from panel E across TCOF1 dosage conditions. n=3 independent differentiations. Dunnett’s multiple comparisons test. **(G)** Representative images of wild-type CNCC differentiation at day 1 and day 4 showing uniform neuroectoderm sphere formation. Scale bars, 1000 μm.

**Figure S2.**
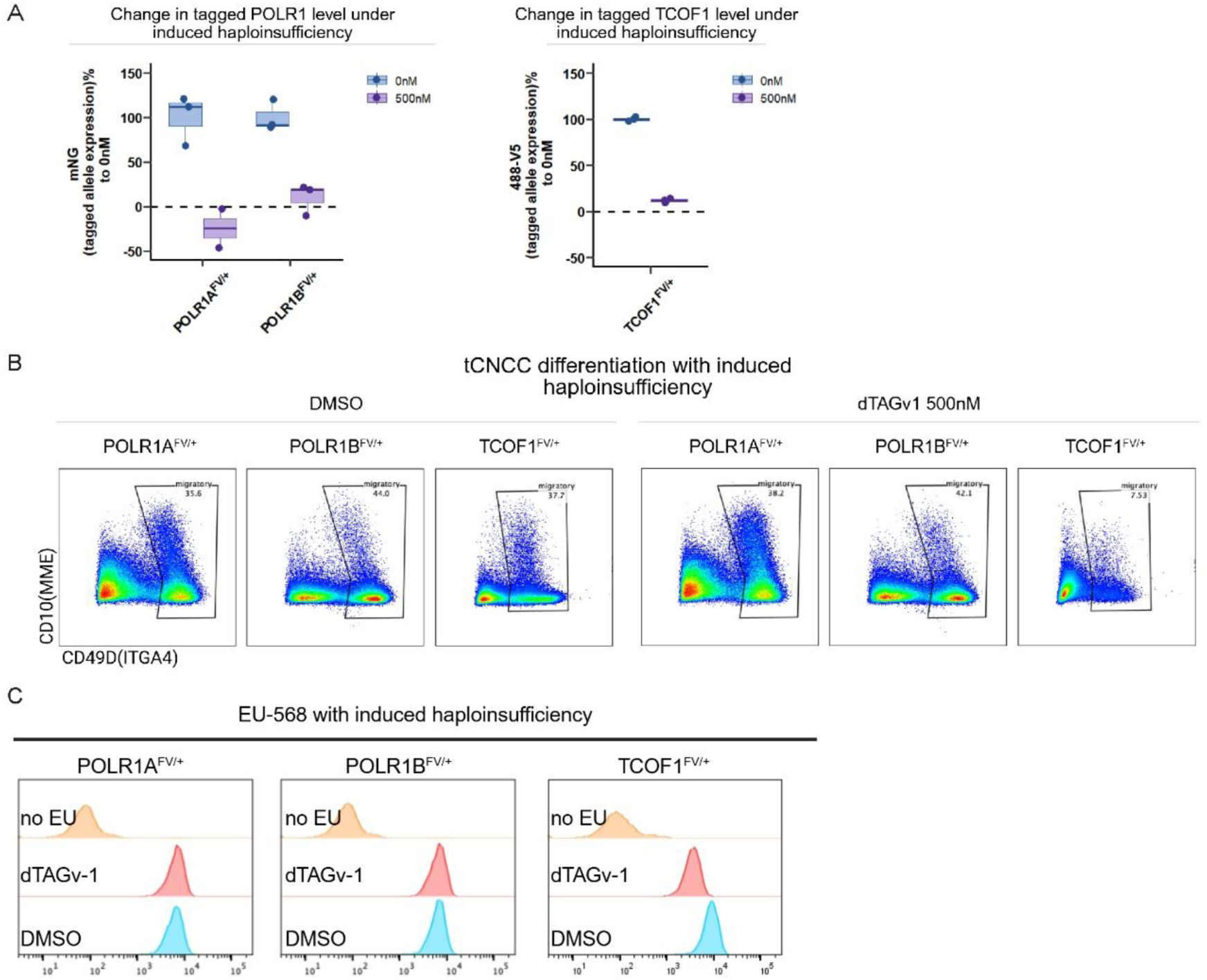
The distinct TCOF1 dosage sensitivity. **(A)** Quantitative evaluation of induced haploinsufficiency (24 hours dTAGv-1 treatment) of *TCOF1, POLR1A,* or *POLR1B* in tCNCC3, as % of remaining tagged factor (product of tagged allele) relative to vehicle treated control cells. measured by flow cytometer of mNeonGreen fluorescence (POLR1A, POLR1B) or 488-V5 staining (TCOF1). n=3 independent differentiations. Statistical comparisons control(0nM) vs dTAGv-1(500nM): POLR1A (p = 4.2×10^-3^), POLR1B (p = 3.2×10^-3^), TCOF1 (p = 1.9×10^-6^), two sided t-test. **(B)** Representative flow cytometry plots showing tCNCC yield (measured by surface marker staining and flow cytometry) following induced haploinsufficiency of *TCOF1, POLR1A,* or *POLR1B*. **(C)** Representative flow cytometry plots showing rRNA synthesis(5-EU incorporation) in tCNCC3 following induced haploinsufficiency of *TCOF1, POLR1A,* or *POLR1B*, measured by flow cytometer of incorporated 5-EU with click-568 fluorescence.

**Figure S3.**
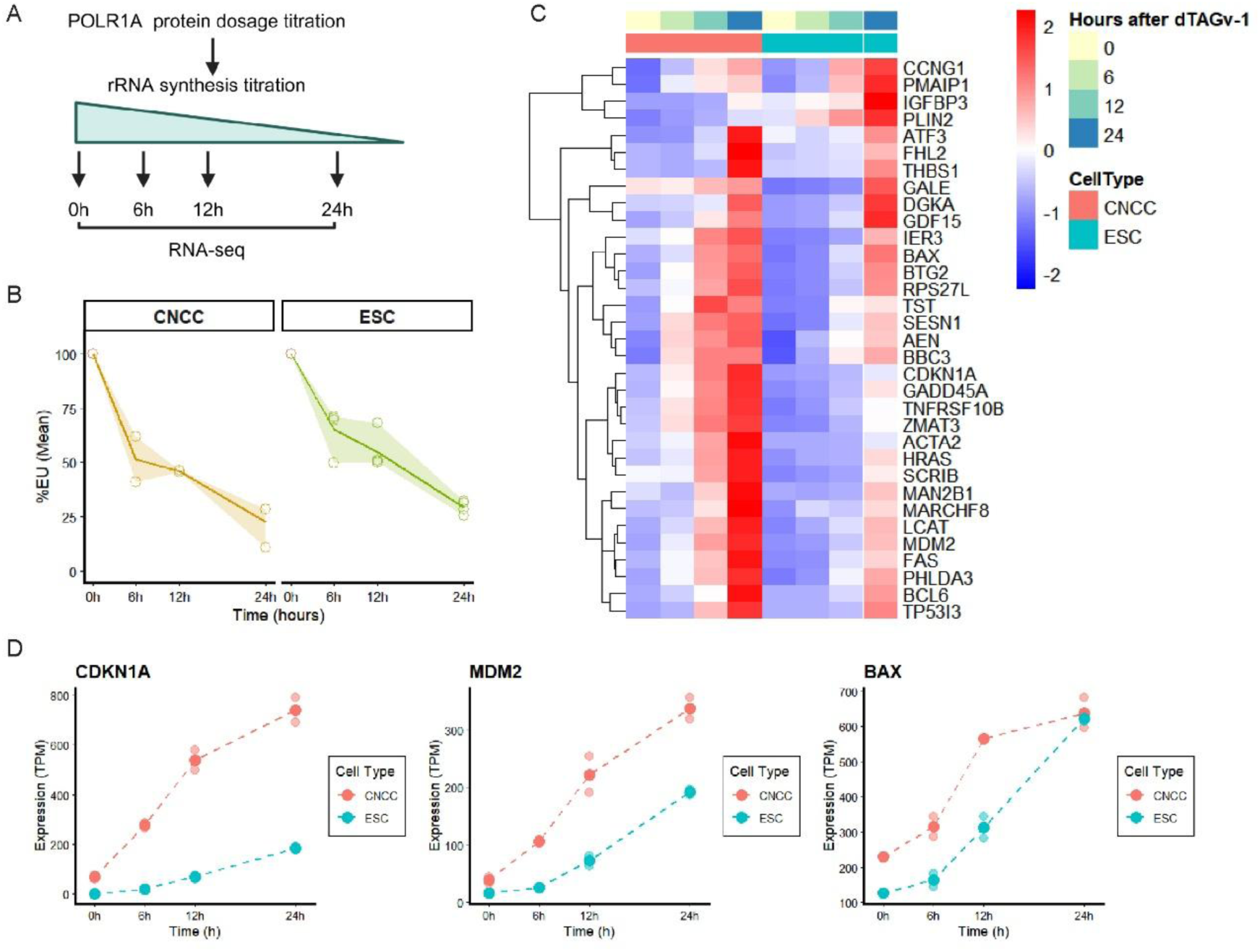
The CNCC-specific vulnerability to rRNA synthesis reduction. **(A)** Schematic overview of titrating rRNA synthesis followed by transcriptome profiling (RNA-seq) in CNCC and ESC. **(B)** rRNA synthesis (5-EU% relative to 0h control) in CNCC and ESC, demonstrating matched rRNA synthesis reduction between the two cell types at each time point. **(C)** Heatmap showing expression dynamics of nucleolar stress response genes in CNCC and ESC across rRNA synthesis titration, revealing cell type-specific sensitivity to rRNA synthesis perturbation. **(D)** Representative expression trajectories of individual nucleolar stress response genes from panel C, highlighting divergent responses between CNCC and ESC to matched rRNA synthesis reduction.

**Figure S4.**
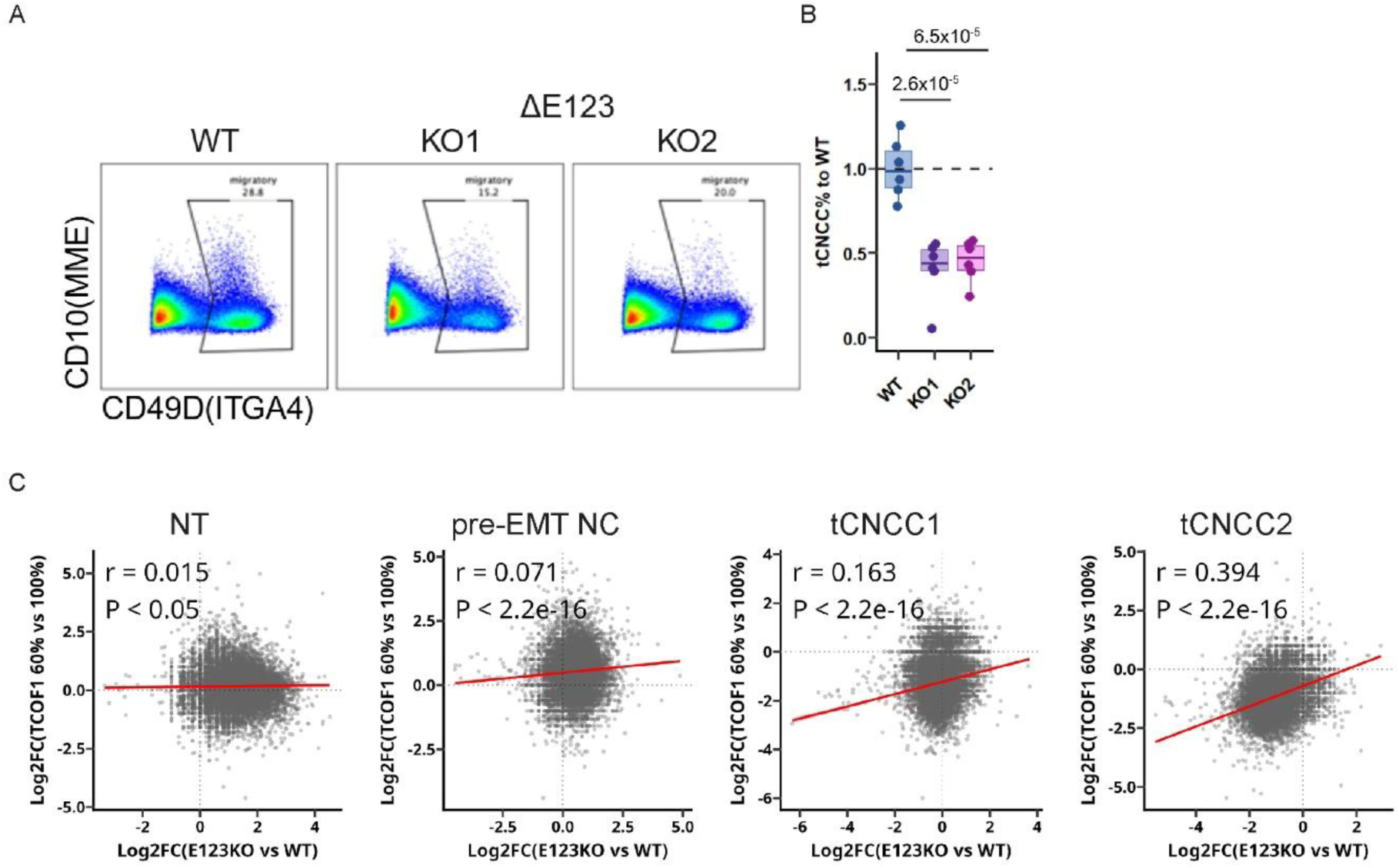
*TCOF1* enhancers E1/2/3 are required for CNCC differentiation. **(A)** Representative flow cytometry plots showing tCNCC yield (measured by surface marker staining and flow cytometry) in wild type and two E123 knockout clones. **(B)** Quantification of tCNCC percentage from panel A. n=6 independent differentiations. Dunnett’s multiple comparisons test. **(C)** Pearson correlation analysis comparing transcriptome-wide expression changes between E123KO vs. WT and TCOF1-depleted (60% protein) vs. control (100% protein) in NT (n=24850 genes), pre-EMT NC (n=24257 genes), tCNCC1 (n=24543 genes), tCNCC2 (n=22567 genes). Each dot represents one gene. Log₂ fold-changes were calculated using DESeq2 from 2 independent differentiations per condition. P-value from t-distribution (H₀: ρ=0, df=n-2).

**Figure S5.**
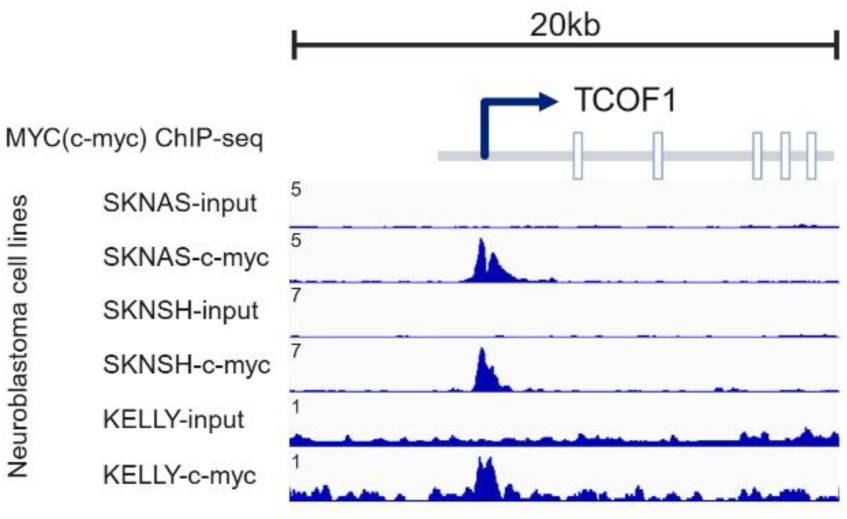
MYC binds directly to the *TCOF1* promoter. MYC (c-myc) ChIP-seq tracks from neuroblastoma (neural crest-derived cancer) cell lines at the *TCOF1* locus, showing robust MYC occupancy at the TCOF1 promoter. Data from Upton et al. 2020, GSE138315.

**Table S1.**
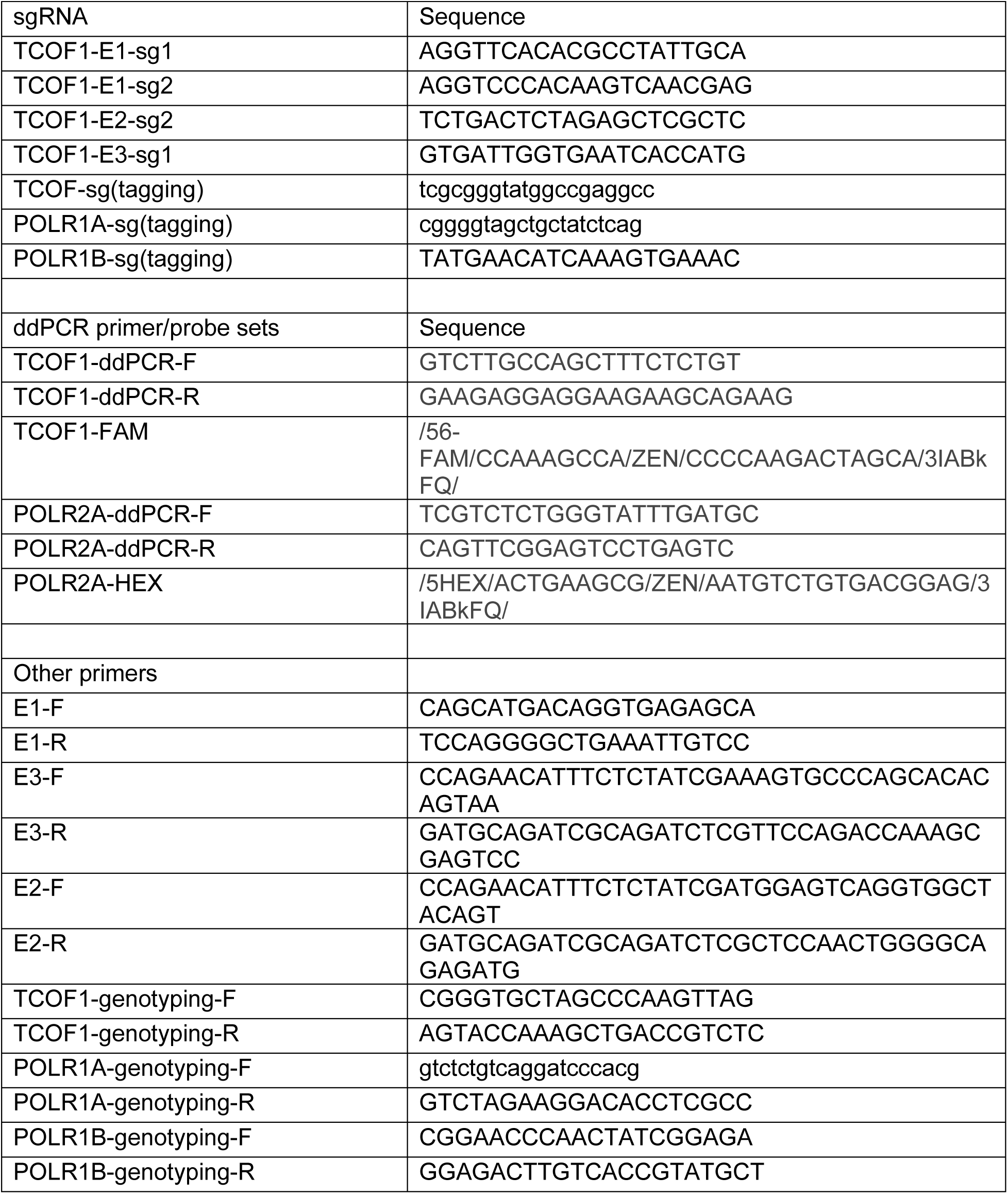
Primer, probe, sgRNA.

**Table S2.** GEO Accession numbers of publicly available datasets.

| Accession | Samples and assays | Reference |
| --- | --- | --- |
| GSE230319 | CNCC ChIP-TWIST1, ChIP-H3K27ac(control and TWIST1 depleted), ATAC(control and TWIST1 depleted) | S. Kim et al, 2024 |
| GSE145327 | H9 hESC, CNCC ATAC | H. K. Long et al, 2020 |
| GSE70751 | CNCC ChIP-H3K27ac, H3K4me3, CTCF, RAD21 | Sara L. Prescott et al, 2015 |
| GSE138315 | neuroblastoma cell lines SKNAS, SKNSH, KELLY ChIP-MYC(c-myc) | K. Upton et al, 2020 |

Data S1. Dosage sensitivity and variance of genes

Data S2. DSH gene features

## Notes

### Competing Interest Statement

The authors have declared no competing interest.

### Summary of Updates

This publication was given a CC-BY-NC-ND license. HHMI's Immediate Access to Research policy requires that preprints that represent major contributions from HHMI labs be made publicly available under a CC BY 4.0 license to enable maximal accessibility, reuse, and distribution of the discovery with appropriate attribution.

